# Tobacco twin and single spots result from genome instability, chromosome breaks, and translocations

**DOI:** 10.1101/2025.11.09.687472

**Authors:** Isabelle J. DeMarco, Kirk R. Amundson, Angelina Phan, Isabelle M. Henry, Luca Comai

**Affiliations:** Department of Plant Biology and Genome Center, University of California Davis, Davis, California 95616, USA; department of Biology, University of Massachusetts Amherst, Amherst, Mass., USA

## Abstract

Somatic recombination can have profound consequences including clonal evolution. In plants, however, the contribution of homologous recombination (HR) to genomic instability remains unclear. The semidominant *sulfur* mutant of tobacco, named for its yellow foliage resulting from a chlorophyll defect, spontaneously develops green and white twin spots that have been attributed to mitotic crossovers. To test this hypothesis, we sequenced DNA from twin spots and surrounding tissues in *su/+* F1 hybrids, expecting somatic crossovers to result in reciprocal chromosome arm exchange in the paired sectors. Contrary to expectations, 20 of 22 twin spots exhibited aneuploidy or reciprocal translocations between non-homologous chromosomes. One twin spot showed a pattern consistent with a reciprocal crossover. Single white or green spots were ∼10X more frequent than twin spots. Among 26 single spots, 18 resulted from terminal arm deletion, 2 from aneuploidy, and 4 were associated with translocations. Deletion and reciprocal exchange breakpoints clustered near the centromere. Together, these findings indicate that nearly all spots arise from deletions, translocations, and other rearrangements likely mediated by chromosome missegregation or nonhomologous repair rather than by HR. These rearrangements altered the ratio of mutant to wild-type alleles and consequently chlorophyll levels. We conclude that mitotic recombination between homologous chromosomes is rare, and that genome instability is the dominant driver of somatic chimerism in this system.

## Introduction

Recombination in somatic cells is a DNA repair mechanism used to repair spontaneous and induced dsDNA breaks. The type of repair determines the outcome. Ideally, this entails restoration of the original genomic status. Often, however, repair results in inherited changes that can affect the phenotype. In heterozygous organisms, an undesirable outcome is Loss of Heterozygosity (LOH); for example, when a *+/m* cell becomes *m/m* or *m* (where *+* and *m* represent, respectively, WT and mutant allele). During development, cells affected by LOH form mutant sectors. In plants, these can become clonal variants, a phenomenon important in plant breeding (Foster and Aranzana 2018). In humans, LOH sectors can form malignant cancers (Happle 1999). The mechanisms underlying these phenomena are of basic and applied importance, affecting, for example, the application of CRISPR to plant genome editing and directing specific recombination outcomes (Guzmán-Benito et al. 2023)

An intriguing manifestation of LOH are twin spots, adjacent epidermal or colony patches that display distinct traits on a different background. They can be found in yeast, insects, and plants. In Drosophila, for example, singed and yellow twin spots can result from a double repulsion heterozygote (Stern 1936) such as *singed* + / + *yellow* if a G2 crossover (CO) occurs between the centromere and the linked *singed* and *yellow* loci. Segregation of the resulting recombinant chromatids can form a spot homozygous for *singed (singed +* / *singed +)* and an adjacent spot homozygous for *yellow (+ yellow / + yellow)* (Stern 1936). Incompletely dominant alleles at a single locus also enable tracing mitotic crossovers as twin spots (Fig. 1, Suppl. Fig. S1). One such locus, *sulfur (su)*, was used in tobacco to demonstrate mitotic recombination by the appearance of green (presumed +/+) and white (presumed *su/su*) twin spots on a yellow-green background (*su/+*) (Carlson 1974). In arabidopsis, soybean, and tobacco, twin spots have been equated to mitotic crossover (Vig 1973; Barrow et al. 1973; Christianson 1975; Mericle and Mericle 1973; Gorbunova et al. 2000; Lörz and Scowcroft 1983; Dulieu 1975; Harrison and Carpenter 1977; Evans and Paddock 1976; Hirono and Redei 1965) (Suppl. Fig. 1-A,B). In summary, considering an *m/+* heterozygote, a reciprocal crossover event exchanges sections of two homologous chromatids: one carrying *m,* the other *+*. The resulting daughter cells inherit *+/+* or *m/m* complements (Fig. S1). This mitotic event would have the same genomic consequences of a meiotic crossover.

**Figure 1.**
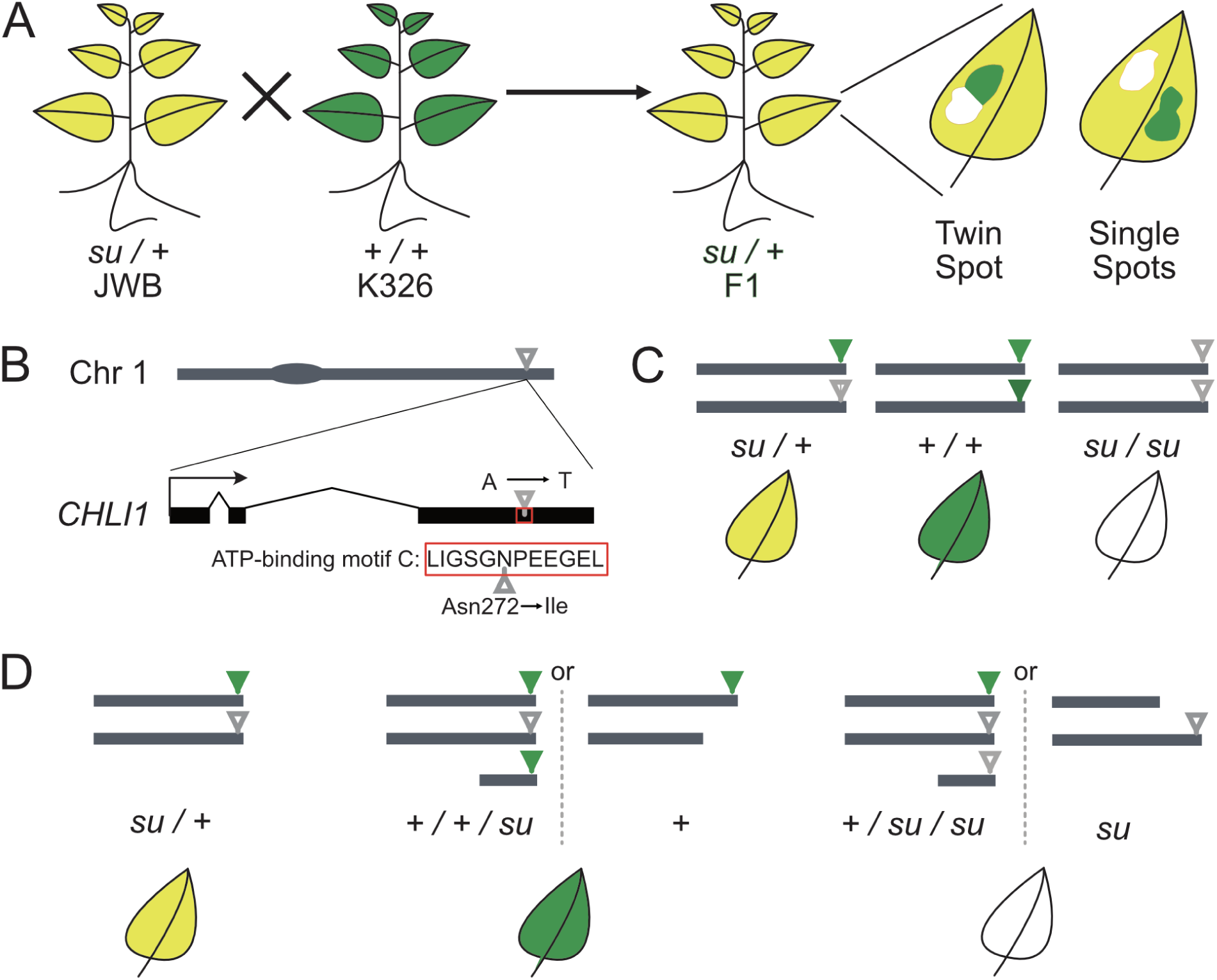
Experimental design and expectations. (A) The plants used in this experiment were F1’s of the *sulfur* heterozygote JWB and of the reference genome variety K326. Leaf color spots due to variation in chlorophyll content arose spontaneously when the *su/+* plants were grown in growth chambers under optimal conditions. (B) Location of the *CHLI1* locus on Chr 1, with the causal SNP (grey lollipop) in the *CHLI1* gene model affecting the indispensable N residue in the predicted ATP binding motif (Walker and Willows 1997). (C) Predicted genotype of somatic variant tissues produced by mitotic crossovers, i.e. homologous recombination between chromatids of different homologs according to Carlson (1974). (D) Alternative hypothesis involving allele dosage change where the ratio of *su* and *+* alleles results in differential tissue colors.

Repair of a dsDNA break, however, can produce twin spots by mechanisms other than reciprocal homologous COs. For example, rearrangements caused by transposon-triggered instability (Dooner and Belachew 1989; Peterson and Yoder 1993) can yield daughter cells carrying, respectively, *+/+/m* (green) and *m* (white), prompting similar explanations for unknown twin-spot syndromes (Zhu et al. 1995). Simple spots, which often accompany twin-spots, are best explained by instability resulting in deletion or other abnormalities (Vig 1973). Intriguingly, syndromes with frequent twin-spotting can arise after genetic transformation (Gorbunova et al. 2000; Zhu et al. 1995) raising the possibility that persisting genomic instability, which can occur as a result of somaclonal variation (Fossi et al. 2019), underlies twin-spot formation.

To evaluate the molecular basis of twin spots, we chose a well characterized system: the incompletely dominant *sulfur* mutation of tobacco (*Nicotiana tabacum*). Tobacco *su/+* heterozygotes are yellow-green due to insufficiency of Mg Chelatase, a multi-subunit enzyme necessary for formation of chlorophyll (Hansson et al. 2002). Incomplete dominance is thought to result from complex poisoning by the mutant subunit (Fitzmaurice et al. 1999; Papenbrock et al. 2000). By analyzing the dosage and sequence of multiple twin and single spots, we show that the majority of twin spots are not the result of mitotic crossover between homologous chromosomes. Instead, they result from the combined effect of genomic instability and dosage effects involving one or the other of two homoeologous *CHLI* loci. The analysis of single spots confirmed the frequent instability affecting these loci. This finding underscores the rarity of homologous recombination in the plant soma, highlighting the frequency of genome instability during leaf development.

## Results

### Experimental design and identification of the *sulfur* mutation

The *sulfur* mutant likely derives from a mutation in the *CHLI1* gene, which has been located to chromosome 1 via transposon tagging (Fitzmaurice et al. 1999) and encodes a subunit of the magnesium chelatase protein complex (Walker and Willows 1997). However, the causal mutation remains unknown, hindering the genotyping of single and twin spots. To identify the causal mutation, we amplified and sequenced the coding sequence of the candidate *CHLI1* locus from leaf genomic DNA of *su/+* and WT siblings in a homozygous John Williams Broadleaf (JWB) background. In half of cloned PCR products from *su/+*, we detected a heterozygous SNP in the third exon that was absent from the WT sibling. This SNP results in an N272 to I substitution (Fig. 1) in the highly conserved ATP binding motif C (LIGSGNPEEGEL) where the mutated N is indispensable (Wang et al. 2023; Walker and Willows 1997). To determine whether this mutant *CHLI1* allele is expressed, we generated and analyzed RNA-seq from individual leaves of six *su/+* and six WT siblings in a homozygous JWB background. Four of six *su/+* samples exhibited heterozygosity of the missense mutation (data not shown). These results, combined with previous transposon tagging, indicates that the *sulfur* phenotype results from a missense mutation in the *CHLI1* coding sequence. The semidominant phenotype of *sulfur* is due to the relative dosage of functional and compromised subunits (Campbell et al. 2014; Hansson et al. 2002; Papenbrock et al. 2000), suggesting that mutations affecting any *CHLI1* homolog may also impart twin- or single-spots. Allotetraploid *N. tabacum* carries two additional *CHLI1* copies on chromosomes 16 and 23. In our leaf RNA-seq data, both genes were expressed, appeared non polymorphic between the two varieties, and lacked any potentially deleterious mutations, suggesting both loci contribute functional CHLI1 protein (data not shown). Therefore, we expected single- and twin-spots would mark copy-neutral loss of heterozygosity on chromosome 1, and changes to dosage of *CHLI1* homeologs on chromosomes 1 (*CHLI1-A*), 16 (*CHLI1-C*) and 23 (*CHLI1-B*), as each outcome is expected to alter the dosage of functional and compromised CHLI1 proteins.

To track mitotic recombination breakpoints, we sequenced twin spot sectors in an F1 hybrid background. We crossed *su/+* JWB to the reference genome cultivar K326 and obtained F1 progeny segregating 1:1 green: yellow *su*/+ (37 green: 41 yellow: 0 white; P of χ square test for 1:1 hypothesis = 0.65). For sequencing and spot observations, offspring of a cross between the above mentioned su/+ JWB and wildtype cultivar K326 were used (Fig. 1A). K326 was selected as the most recent reference genome by Sierro et al. was made with this cultivar. We can distinguish homologues in the hybrids using extant SNPs between the two parents, allowing for detection of loss-of-heterozygosity events.

### The chimeric patterns of *sulfur* spotting are often clustered and layered

Upon selfing, the *su/+* heterozygotes produced progeny in the expected Mendelian 1:2:1 ratio corresponding to *su/su*, *su/+*, and *+/+* genotypes. Albino *su/su* individuals were unviable and only detectable when germinated on agar medium. On soil, the progeny exhibited a 2:1 ratio of heterozygotes to WT plants. Heterozygous seedlings were distinctly yellow and slower-growing than their WT siblings, but gradually became larger and greener (Fig. 1A). These *sulfur* heterozygotes in the JWB background were crossed with WT cultivar K326. On both soil or agar, the resulting seed yielded progeny displaying the expected phenotypic ratio of 1:1, green:yellow. The yellow *su/+* F1 hybrids were used throughout this work. Their genome-wide heterozygosity enabled haplotype identification through SNP analysis.

We quantified mosaic sectors in 25 *su/+* hybrids grown under optimal controlled environmental conditions. For clarity, we refer to the background leaf color as yellow, and to the sectors as green or white, although their shades varied. Spots appeared spontaneously on developing leaves throughout growth (Fig. 2C–E), affecting all regions of the blade, from center to margin. Although often located near midribs and major veins, they typically did not span across them. All 25 plants exhibited at least one green and one white single spot (GSS, WSS, respectively), and all but two also had at least one twin spot (TS) (Fig. 2F). Green single spots were most common, followed by white single spots and then twin spots, occurring at an approximate 18:4:1 (GSS:WSS:TS). The deficit of white spots could be due to the difficulty in detecting them. Most spots measured between 1 and 5 mm in diameter. According to clonal analysis, the events generating them occurred around the 3mm primordial leaf stage (Poethig and Sussex 1985a).

**Figure 2.**
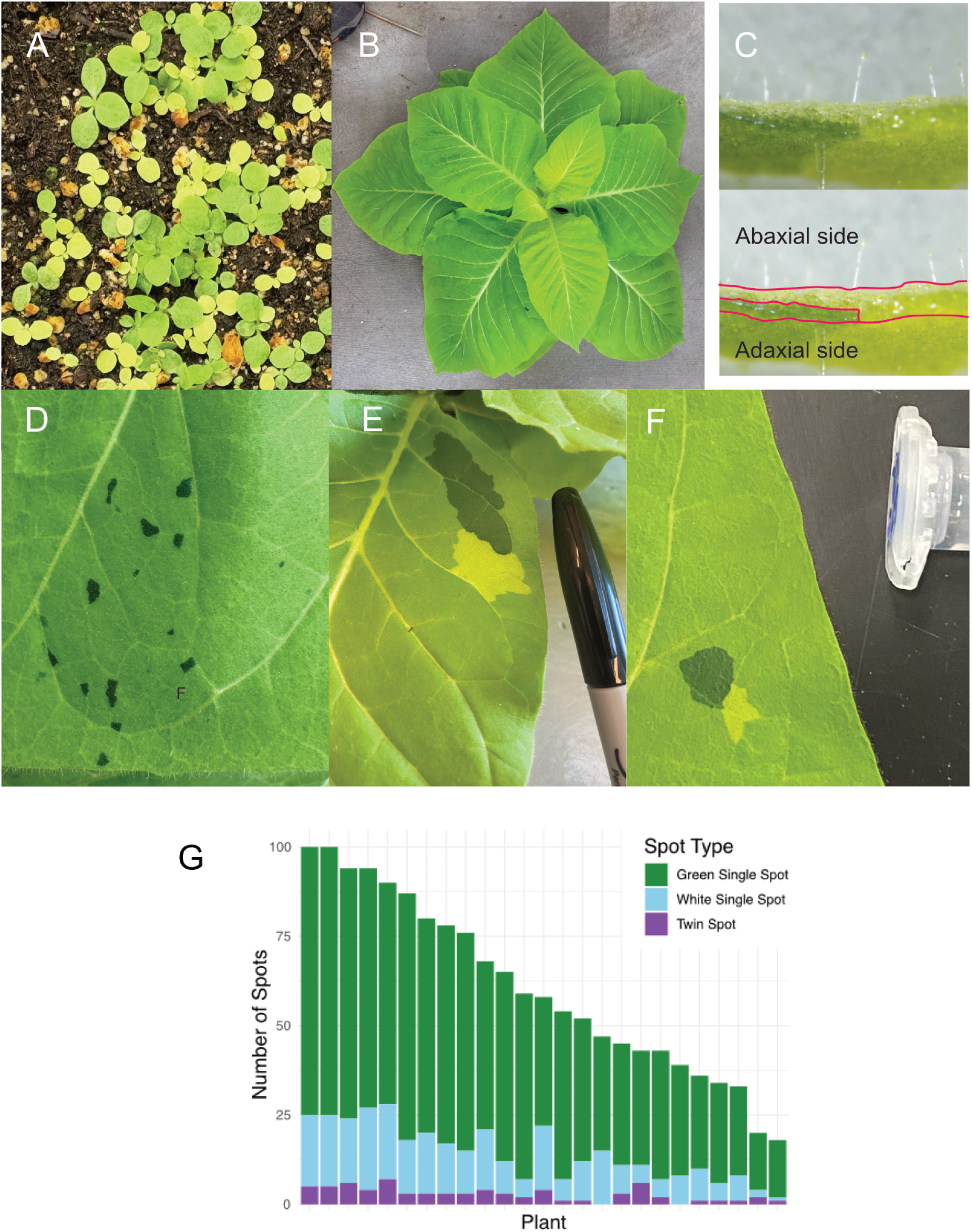
Manifestation of spotting in *sulfur N. tabacum.* A) Young F1’s of a *su/+* X *+/+* cross showing 1:1 trait segregation, approximately 2 weeks after planting. B) A *sulfur* heterozygote at 12 weeks of age. Notably, as *su/+* plants mature, they become larger, more vigorous, and the leaf color shifts from yellow to pale green. C) Cross-section of a green spotted leaf, showing the layer-specificity of the green cells. The margins of the cut and the green spot cross-section are traced in the bottom image. The adaxial leaf surface is down. D) A cluster of green spots on a leaf. The majority of spots were <5mm in diameter. E, F) Twin spots. G) Count of green spots, white spots, and twin spots >1 mm in mature leaves of 25 individuals sampled when they displayed ∼5 mature leaves (3-8 range).

Among ∼1500 total spots, only 26 green single spots, 7 white single spots, and 11 twin spots ranged from 5 mm to 1 cm, while 6 green single spots and 1 white spot exceeded 1 cm (Suppl. Table S1). Increased twin spot size could of course depend on their intrinsically duplicated nature. While these data reflect a subset of 25 plants, over 200 plants were examined throughout the study in both growth chambers and greenhouses. In one instance, we observed a green sector spanning an entire leaf indicating that a mutant cell generated the whole primordium (Poethig and Sussex 1985b). Additionally, many spots smaller than 1mm were observed on leaves displaying larger spots, but were not counted due to the difficulty of reliably detecting white single spots and twin spots of this size.

We observed additional features of spot formation. Many spots exhibited clear cell-layer chimerism and appeared on a single leaf surface—most frequently the adaxial. Their layer arrangement affected intensity and visibility (Suppl. Fig. S2): green spots on the abaxial surface were only faintly visible from the adaxial side. Cross-sectional analysis confirmed the presence of chimeric layers (Fig. 2C). For example, a sampled green sector did not extend through the entire leaf thickness, but was limited to the palisade and parts of the spongy mesophyll. In two cases, a twin spot displayed break-down of a spot in a mosaic of tiny patches (Suppl. Fig. S3).

Clustering of same-colored spots was relatively common (Fig. 2D, Suppl. Fig. S4), and high-magnification inspection revealed green microspots surrounding larger sectors (Suppl. Fig. S5). Although similar small white spots may exist, they would be harder to detect. In conclusion, single spots were the most frequent and there was variation in the number of spots between plants and between leaves of the same plant (Suppl. Table S1). Twin spots were less common, and microspots appeared frequently associated with larger events.

### Most Twin Spots display reciprocal dosage changes

We aimed to determine the molecular recombination mechanism underlying the formation of twin and single spots in *sulfur* tobacco. Specifically for twin spots, we sought to distinguish between homology-dependent recombination resulting in reciprocal crossovers and more chaotic events such as chromosomal translocations by the connected genomic predictions (Fig. 1, Suppl. Fig. S1). To be able to simultaneously detect copy number variation and Loss-Of–Heterozygozity, we leveraged the extensive polymorphism between cultivar JWB (which produced the original *sulfur* mutation) and cultivar K326. We then sequenced genomic DNA isolated from 28 single and 22 twin spots arising in yellow *su/+* F1 offspring.

We sequenced DNA from a total of 50 spot events covering 22 twin spots, 17 green single spots, and 11 white single spots. For each twin spot pair, we sequenced DNA from the green, white, and the adjacent yellow background. For single spots, we sequenced DNA from the spot and the adjacent background. The number of reads across the genome was standardized relative to a control sample, and plots were analyzed for regions of differential coverage (Fig. 3A, C), to assess chromosomal copy number. Yellow tissue served as euploid control where all chromosomes should display 2 copies. This assumption proved correct in all cases except one where a control sample displayed a terminal arm deletion that did not involve Chr 1 or Chr 23 and therefore did not result in an altered color phenotype (Fig. 3C).

**Figure 3.**
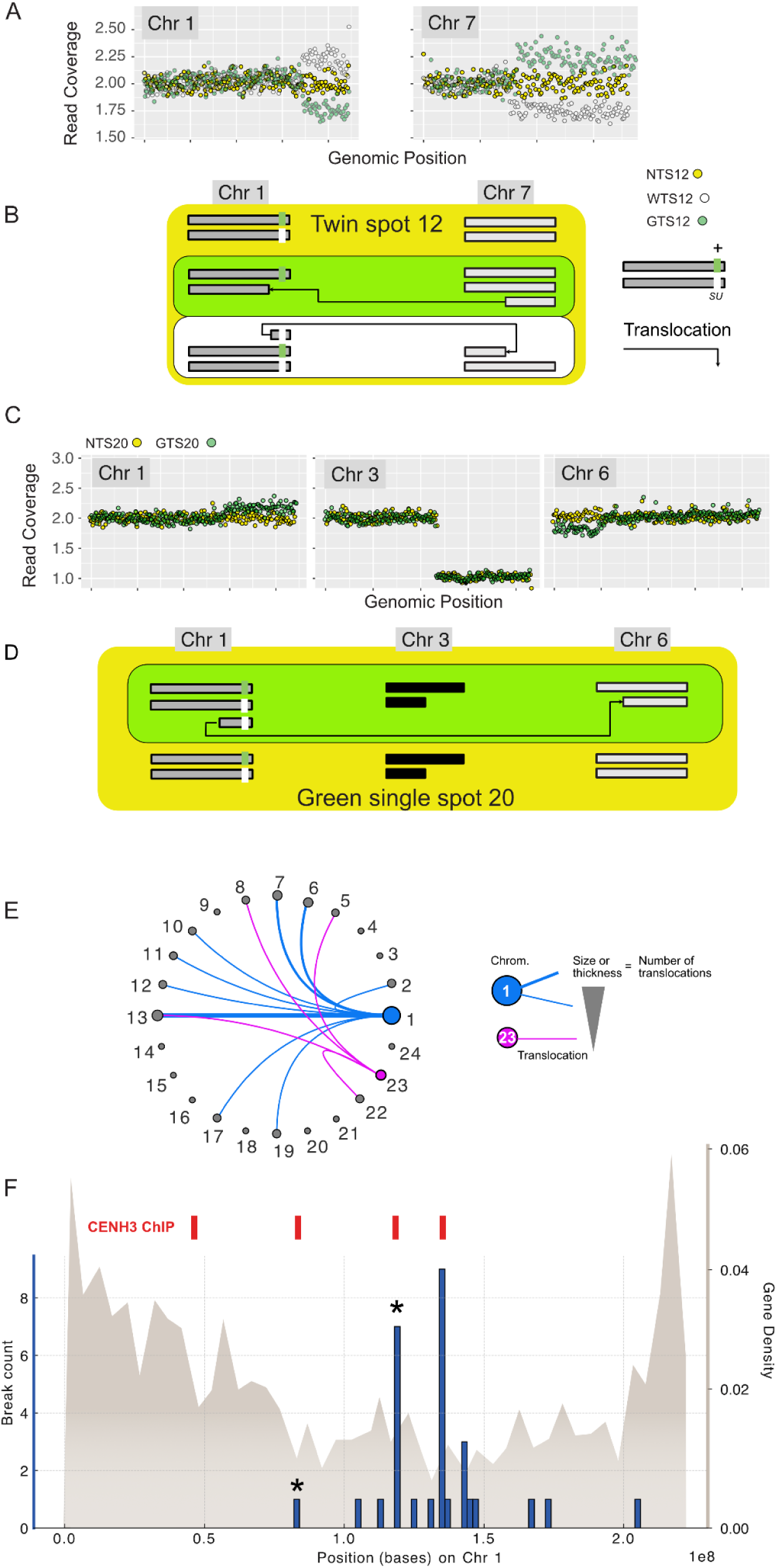
Genomic analysis of DNA in leaf spots. (A,B, and C) Sequencing read dosage analysis (top) and karyotypic interpretation (bottom). Each dot represents the 2-copy standardized dosage for a 1Mb window (see Methods). Segments with dots in the 3X range are considered duplications. Segments with dots in the 1X range are considered deletions. The spots, however, are frequently chimeric (Fig.1D) resulting in dilution of green or white cell DNA with DNA from background yellow cells, resulting in a dosage change proportional to the dilution. For example, if DNA carrying a duplication is diluted 1:1, dosage will shift to ∼2.5 instead of 3.0. (A). Twin spot 12 (TS12) DNA from the yellow-green background tissue (NTS12), from the green spot (GTS12), and from the adjacent white spot (WTS12). In the twin spot DNA, reciprocal segmental dosage changes are apparent for Chr 1 and Chr 7. (B) Representation of the karyotype of background, white and green tissue. (C) Analysis of DNA in green single spot 20 (GSS20). NGSS20: background tissue from the leaf that formed green single spot 20. The dosage change displayed on Chr 3 indicates a deletion not involved in the color spotting trait. The green spot displays duplication of the distal right arm of Chr 1 and a concurrent deficiency of a distal left arm of Chr 6. This is likely a translocation between the two arms. The stripes represent the *CHLI1* locus and its color represents the WT (+, green) and mutant (*su*, white) alleles, respectively. There is a large difference in the dosage shift displayed by Chr 1 and 6 (reduced representation of the affected layer) in comparison to Chr 3 (somatically fixed). (D) Representation of the karyotype of background and green tissue (C) Translocation partners for Chr 1 and Chr 23. (D) Break point analysis of Chr 1 with instances plotted in the context of gene density and the reported positions of CENH3-enrichment (Zan et al. 2025). The position of the two breaks associated with isochromosomes formation in TS23 (Suppl. Fig. S6) are marked (*).

We considered two contrasting hypotheses: precise homologous recombination affecting the *CHL1-A* gene on Chr 1 should be copy number neutral or, alternatively, dosage changes affecting the *CHLI1* genes, either *CHL1-A* on Chr 1 or *CHL1-B* on Chr 23 would alter the balance of mutant and WT *CHLI1* alleles and potentially explain differences in chlorophyll production.

The twin spots displayed a range of genome instability outcomes. With the exception of twin spots 20 (poor sequencing data) and 27 (discussed later), all twin spots displayed dosage shifts in Chr 1 or 23, but never both. In one case, twin spot 39, the dosage change was not symmetrical: Chr 23 was monosomic in the white spot, but, surprisingly, also in the green spot (Suppl. Fig. S2). It was classified as undetermined. In the 19 remaining twin spots, the DNA from the green and white twin sectors displayed symmetrical and reciprocal dosage changes. In four cases, these changes affected a whole chromosome carrying a *CHLI1* locus, either 1 (three cases) or 23 (one case) corresponding to trisomies where one of the twins lost and the other gained an entire chromosome. In fourteen twin spots, the chromosomal arm carrying *CHLI1* (10 *CHL1-A* on Chr 1 and 4 *CHLIB* on Chr 23) had undergone a reciprocal translocation with another chromosome, which, in most, cases appeared randomly chosen (Table 1, Fig. 3E). The one remaining twin spot displayed green-white reciprocal changes that were limited to one or the other arm of Chr 1. The white spot had three copies of the *CHL1A* gene, the green spot a single copy. This is consistent with two breaks leading to symmetrical formation of two isochromosomes (Suppl. Fig. S6). In conclusion, the analysis indicated that, in 19/22 twin spots, color variation could be explained by chromosome reciprocal dosage variation of a *CHLI1* allele on either Chr 1 or Chr 23, either translocation or aneuploidy (Table 1).

**Table 1:**
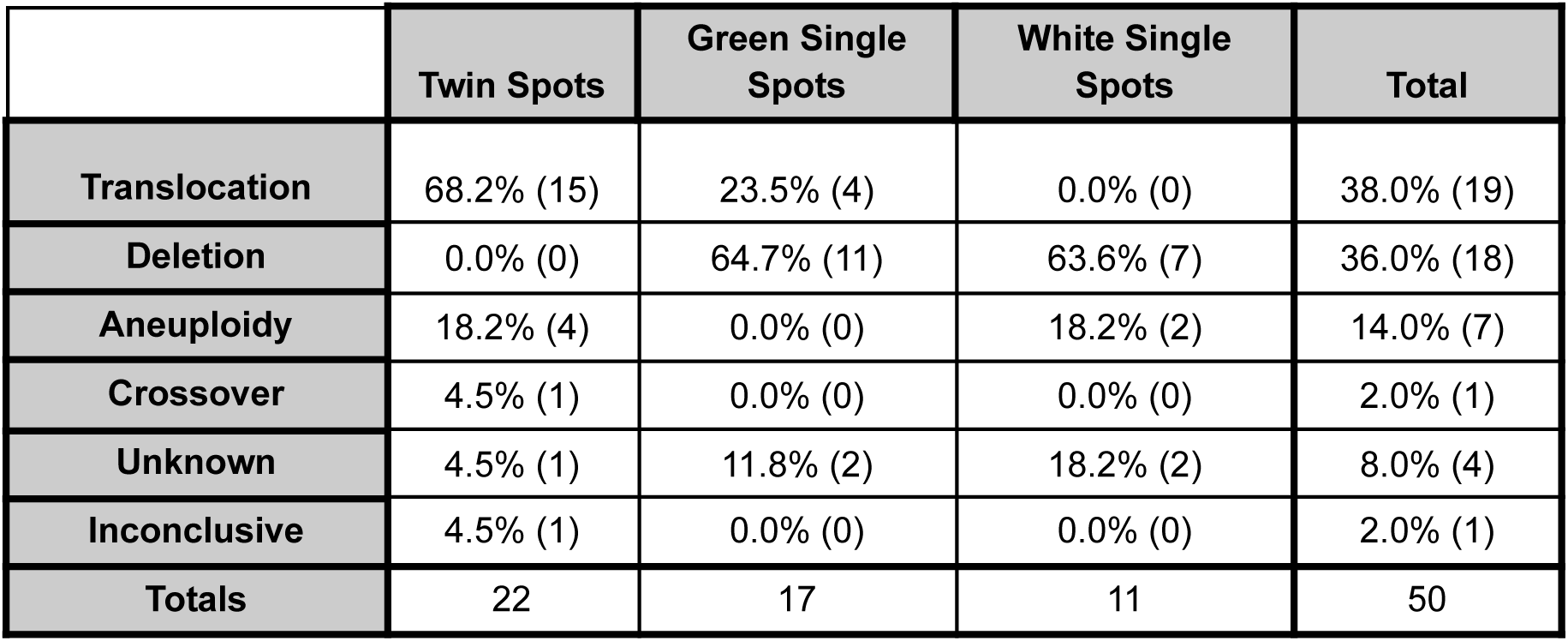
Genome changes infrared in each spot type from dosage plots.

### Single spots derive from deletion, translocation, and aneuploidy

The analysis of 28 single spots, 17 green and 11 white, identified dosage changes in *CHLI1* in 26 samples, with the remaining 4 displaying apparently normal genomes. *CHLI1* dosage alteration could yield both color types. Three spots had *CHL1-B* on Chr 23 affected, in all cases resulting in white spots associated with the loss of one copy of *CHL1-B*. The remaining 21 cases affected *CHL1-A* on Chr 1, with 6 resulting in white spots, and 15 in green spots (Table 2). The most common change was deletion (18), followed by translocation (4), and aneuploidy (2) (Table 1). Notably, the 4 translocations were only observed in green spots, while the 2 cases of aneuploidy were in white spots. Setting aside the four unresolved cases (see below for further analysis), the rest conformed to the pattern of dosage changes identified for twin spots.

**Table 2:** *CHLI1*-containing chromosome changes in each spot type. (note: does not include spots categorized as potential crossover). For each category, the number of samples is indicated in parentheses.

|  | Twin Spots | Green Single Spots | White Single Spots | Total |
| --- | --- | --- | --- | --- |
| **Chromosome 1** | 71.4% (15) | 100% (15) | 66.7% (6) | 80.0% (36) |
| **Chromosome 23** | 28.6% (6) | 0% (0) | 33.3% (3) | 20.0% (9) |
| **Totals** | 21 | 15 | 9 | 45 |

### Chimerism biases the measurement of dosage changes

Interestingly, the amplitude of the observed dosage changes varied from spot to spot and was always lower than the 1X or 3X levels expected for homogeneous monosomy and trisomy. We deduced that this was caused by layer chimerism through dilution of the mutant DNA by DNA from background cells. In addition to the chimerism detected in leaf sections, two single spots displayed zones with different colors due to differential layer invasion by the mutant cells. In one example (Suppl. Fig. S5), both zones display deletion of the Chr 1 arm carrying the *CHLI su* allele causing increased green coloration. In the light green zone, the right arm of Chr 1 showed a subtle decrease in dosage indicating high dilution. The dosage shift found in DNA from the dark green zone was larger, consistent with an increased fraction of green cells (Suppl. Fig. S5). Therefore, both altered dosage level and color intensity were influenced by layer occupancy by the mutant cells.

### Chlorophyll accumulation is tightly linked to the ratio of *CHL1* mutant and WT alleles

Our analysis is consistent with a dosage model in which the fraction of mutant allele determines the amount of chlorophyll produced. For example, although *CHL1-B,* on Chr 23, is homozygous WT, variation in Chr 23 dosage altered the phenotype by changing the total number of + alleles expressed by the two loci. Specifically, deletion of one copy of *CHL1-B* yielded white tissue with *su/+, +/NA* genotype (2:1 WT/mut ratio). On the other hand, presence of a third copy of *CHL1B* yielded green tissue with *su/+, +/+/+* genotype (4:1 WT/mut ratio). Variation in dosage of the *CHL1-A* alleles on Chr 1 generated similar results but necessitated the ability to distinguish between the WT and mutant alleles. We genotyped the mutation causal SNP and the haplotype of the surrounding region (the K326 haplotype carries the WT allele while the JBW haplotype carries the mutant allele). In all spots with altered Chr 1 dosage we measured the percent reference (+ haplotype of K326) comparing green and white sectors with the yellow background. Fig. 5A reveals a consistent pattern of increased reference (WT) identity in green tissue (K326 haplotype), decreased reference identity in white tissue (JBW haplotype), and ∼50% reference identity in the yellow heterozygous tissue. This trend was also observed in *CHL1-A* amplicon sequencing (Fig. 5B) where green sectors have increased WT/mutant *CHL1-A* allele ratio and white sectors have decreased ratio. Spots with alterations to Chr 23 had the same *CHL1-A* genotype as the heterozygous control tissue, confirming no change in the Chr 1 alleles (Fig. 5B). In conclusion, the combination of genotypic and dosage analysis of the *CHLI* genes indicated that the fraction of *su* allele out of the total resulted in varying chlorophyll content (Suppl. Fig. S7).

### Do dosage neutral events involve mitotic crossovers?

Copy number neutrality of the two copies of Chr 1 in twin spot 27 was consistent with homologous crossover. Such condition could also result from other mechanisms, such as gene conversion and pseudo uniparental disomy, i.e. loss of one homolog coupled to duplication of the other. To distinguish these alternatives, we genotyped Chr 1 by quantifying the mapping efficiency of sequencing reads to the reference genome (see Methods). This provided a quantitative measure for zygosity. In TS27, the right half of Chr 1 diverged from the 50% VAF expected from haplotype analysis and was about 65% in the green sector and 35% in the white sector, indicating LOH consistent with reciprocal crossover (Fig. 4B).

**Figure 4.**
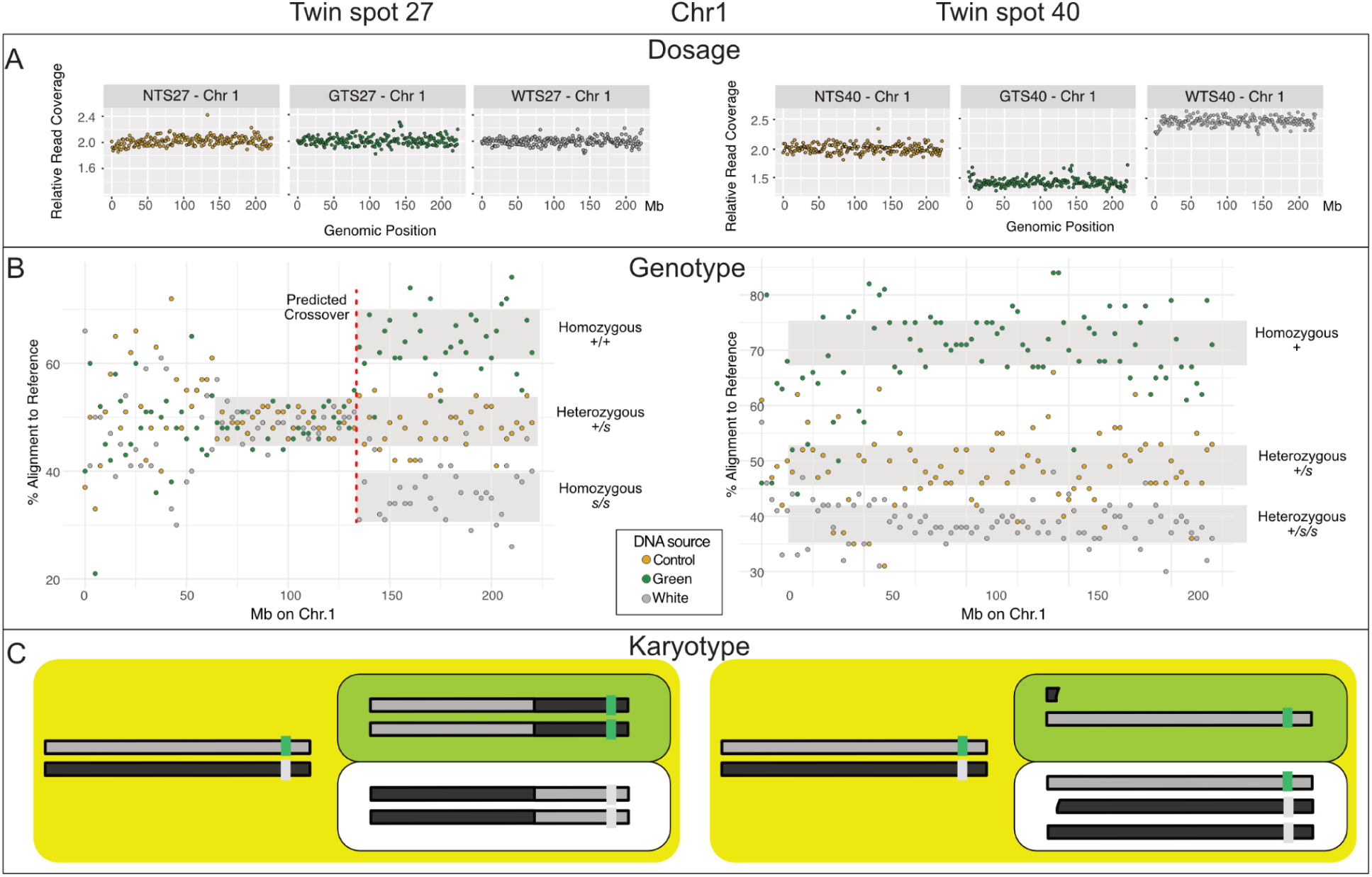
Analysis of dosage and genotype in *sulfur* spots. (A,B) Read dosage plots for twin spots 27 and 40, respectively, demonstrate disomy (A) and aneuploidy (B) for Chr 1. The left tip of Chr 1 might have broken and translocated during missegregation leading to aneuploidy (C,D) Haplotype analysis was performed by determining percent read alignment to the reference sequence of Chr 1, which is the WT haplotype (K326). (C) LOH in the right arm of Chr 1 in the green and white spots vs heterozygosity in background yellow tissue. The red line indicates the position of the predicted crossover. The left chromosome arm is noisy. (D) LOH is visible along the entire chromosome. (E) Percent read alignment to the *CHLI1-A* region of Chr 1 (telomere to break) reference sequence of DNA from green, white, and control yellow tissue. Percent alignment is variable due to layer chimerism, but white spots display low alignments (*su* haplotype) while green spots display high alignment (+ haplotype). (F) Dosage analysis of the causal *su* SNP carried out by Nanopore sequencing of PCR amplicons.

**Figure 5.**
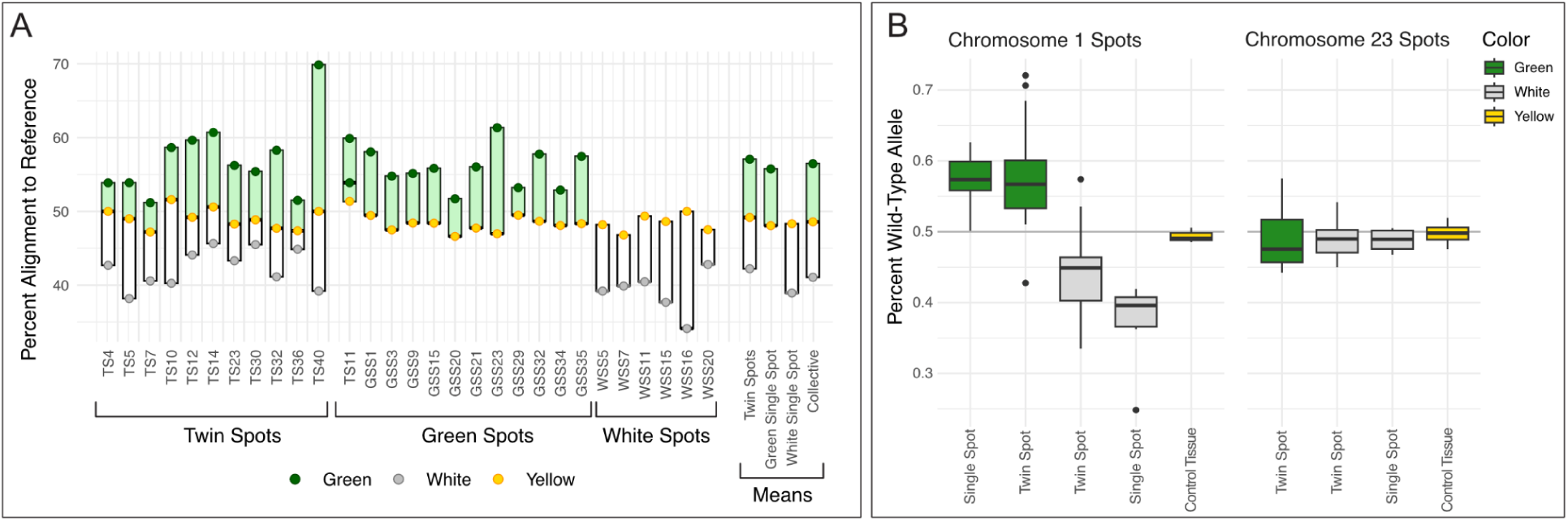
Dosage effects in leaf spots of *sulfur*. (A) Dosage was derived by the percent alignment of sequencing reads to the reference genome. Measurements were binned (see Methods) over the copy-variant chromosomal region carrying the *CHLI1-A* locus. The color spots represent the values obtained for each tissue sample. (B) Dosage derived by counting haplotypes in Nanopore sequenced 2kb amplicons covering the variant SNP in the *CHLI1-A* locus. No copy number variation of the Chr 1 *CHLI1-A* locus was seen for spots derived from Chr 23 *CHLI1-B* locus instability.

**Figure 6.**
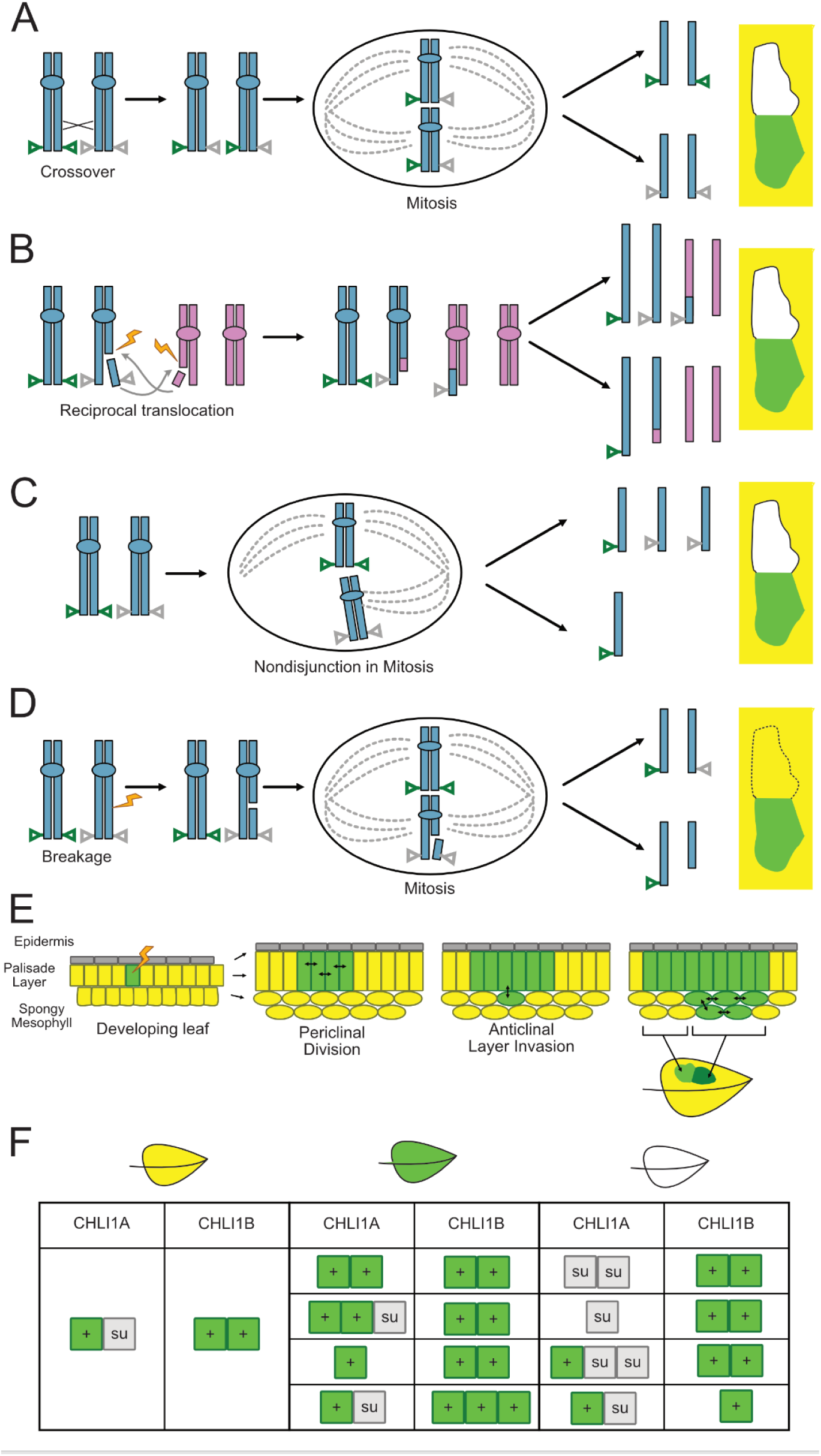
Mechanisms of genome instability and chimeric outcomes. Alternative mechanisms potentially explaining the formation of twin or single spots. (A) Reciprocal crossover between homologous chromosomes followed by partition of the same allele types to daughter cells. (B) Translocation between different chromosomes results in daughter cells with unbalanced dosage of the *CHLI1* region and the translocation partner: 3:1 vs 1:3. (C) Formation of a single spot. Deletion of the distal chromosome arm containing the *CHLI1* gene results in a deficiency. Deletion of the su allele results in a green spot as illustrated. Alternatively, deletion of the + allele results in a white spot (not shown). (E) Most spots involve layer chimerism. At times, layer invasion creates zones in which the mutant cells (green in this case) form regions of different thickness appearing as twin spots with different degrees of green. (F) Allelic interaction at two loci results in yellow, green, or white.

We examined three other events that had genome-wide disomy: 2 green single spots (GSS2, GSS31), and 1 white single spot (WSS10) (data not shown). A fourth spot, WSS1 was included among these candidates because, although it exhibited aneuploidy in chromosomes other than Chr 1 or 23, both *CHLI*-containing chromosomes were disomic (Suppl. Fig. S7). In all cases, we found that heterozygosity was maintained, ruling out chromosome level LOH.

By comparison, haplotype analysis applied to the trisomy-monosomy displayed by twin spot 40 revealed that the green spot cells had lost the chromosome carrying the *su* allele, which in turns had been gained by cells in the white spot (Fig. 4B). In conclusion, we found that Chr 1 right arm LOH explained the white-green twin spot 27, but failed to find Chr 1 LOH for four other single spots that did not display dosage changes. As a result, TS27 was consistent with a reciprocal crossover.

All three of the remaining dosage neutral spots maintained haplotype heterozygosity. For green single spot 31, allelic analysis by amplicon sequencing displayed an increased ratio of WT:mutant allele. This indicated a local change covering the *CHL1A* gene that was consistent with either deletion or gene conversion. The other sequenced spot, white single spot 10, maintained an even ratio of WT:mutant allele. This could be explained by a mutation that inactivated the WT allele, but was not detected in the amplified region. The last two single spots could not be investigated further because all available DNA had been used to construct the sequencing library.

### Chr 1 breaks define fragile sites and the centromere position

The discovery of multiple translocations and deletions affecting Chr 1 provided the opportunity to map the observed breaks (Fig. 3F, Suppl. Table S2). Most breaks occurred between 100 and 150 Mb, with a distinct peak in the 133-135 Mb region. The break-affected region is in the center of the chromosome and displays low gene density, consistent with heterochromatic DNA and a pericentromeric or centromeric nature. According to ChIP analysis (Zan et al. 2025), four chromosomal sites are bound by CENH3 suggesting either a polycentric nature or uneven centromeric function. A majority of breaks appear very close or inside one such CENH3-enriched region. Furthermore, the breaks define fragile sites in the right chromosomal arm and are consistent with centromeric activity left of 105 Mb. Breaks occurring to the left of the centromere will cause deletion or translocation of the left arm and, therefore, should not affect the *CHLI1* dosage. Two breaks that occurred during the formation of isochromosomes in twin spot 23 are very close to the internal CENH3-bound regions. They suggest that the distal CENH3 sites are dispensable for centromeric activity (Suppl. Fig. S6).

Mapping the fragile regions of Chr 23 yielded different results. Our predicted CENH3-binding site places the predicted centromere, indicating it is likely telocentric or subtelocentric, consistent with the conclusions of Sierro et al. (2024). The *CHLI1* homolog on chromosome 23 is located near the opposing telomere. No breaks were observed in or near the predicted CENH3 binding regions. The telocentric nature of chromosome 23 could help explain this observation.

### Accompanying changes indicate common instability

Changes that were presumably coincidental accompanied the instability in the *CHLI1* chromosomes. Four spots displayed extraneous chromosomal changes showing deletion, and aneuploidy events in non-*CHLI* containing chromosomes, indicating presumably independent genome instability events (Fig. 3C, Suppl. Fig. 8). Twin spot 31 displayed trisomy of both chr 1 and 22 in the white sector, while the green sector has low sequence quality. Green single spot 20 displayed translocation of Chr 1 and 6, while Chr 3 was monosomic (Fig. 3C). Monosomy was also detected in the adjacent yellow tissue sample, suggesting that loss of Chr 3 occurred earlier in leaf development. In white single spot 1, no changes to chromosome 1 or 23 were detected, however there was aneuploidy of Chr 20, deletion and duplication in an arm of Chr 3, and uncategorized instability in Chr 6 (Suppl. Fig. S6). Lastly, WSS16 displayed aneuploidy of Chr 1, 11, and 13.

Multiple nonhomologous chromosomes underwent reciprocal translocations with Chr 1 and 23. Twelve different nonhomologous chromosomes were involved in translocations, with 5 being involved in translocation in more than one spot. Chr 13 was the most common translocation partner of Chr 1, occurring in the DNA of three spots and with Chr 23 in the DNA of one spot (Fig. 3E). Three of four breaks clustered in the same genomic bin, which was not a CENH3 enrichment region. There was no observed bias towards translocation with chromosomes of a particular subgenome. When observing the translocation sites among the nonhomologous chromosomes, it was found that the chromosome break occurred in a predicted CENH3 binding region in only 4 of the 19 spots (Suppl. Table S2).

## Discussion

We aimed to elucidate the mechanisms by which chimeric green-white twin sectors are formed on the leaf of heterozygous *sulfur* (*su/+*) tobacco, with a specific focus on evaluating the contribution of reciprocal crossover (CO) events. We determined that the *su* mutation is caused by a SNP in *CHL1-A,* the S-genome copy of *CHLI1.* This SNP causes a missense change that likely inactivates the CHLI1 subunit of Mg Chelatase while preserving its ability to participate in the protein complex. Two homeologs of *CHLI1*, on Chr 1 and one on Chr 23, contribute to the expression of the I subunit. A paralog was detected on Chr 16, but its contribution to Mg Chelatase activity is not clear. In this study, it was not involved in spotting. WT alleles (+) are present on both Chr 1 and Chr 23 and appear equivalent in the spotting assay, whereas the mutant allele resides on Chr 1 and expresses the inactive subunit that poisons the complex. Accordingly, the ratio of WT to *su* alleles determines leaf color phenotype according to the scale shown in Fig. 5 and Suppl. Fig. S9.

Sequencing mosaic sectors arising in *sulfur* heterozygous plants revealed the underlying genomic causes. We detected frequent dosage changes whose amplitude never reached one or three full copies consistent with layer chimerism in all spots. The plant shoot apical meristem (SAM) is organized in three layers, L1, L2, L3, that form, respectively, epidermis, parenchyma (including the photosynthetic palisade and mesophyll cells), and vascular system. Mutations accumulate in a layer-specific manner (Amundson KR, Marimuthu MPA, Nguyen O, Konsam S, Phan A, Demarco IJ, Henry IM, Comai L. 2025; Goel et al. 2024). Twenty of 21 twin spots exhibited changes in allele dosage explaining the color change. Single spots originated predominantly from deletion of the terminal arm of either Chr 1 or 23, or whole chromosome aneuploidy, altering the allelic balance toward green or white. Loss of the *su* allele produced green sectors, while deletion of a WT allele produced white ones. In some cases, the broken chromosomal end translocated to a new genomic location. Most of the numerous breaks in Chr 1 localized to a gene-poor region at its center consistent with the fragility of centromeric regions. The leftmost boundary of observed breaks defines the maximum right extent of the centromere. CENH3 ChIP identified four bound regions on Chr 1 (Sierro et al. 2024; Wang et al. 2024; Zan et al. 2025).

Our data on twin spot 23 suggest that while all the CENH3 sites in this polycentric region may contribute to centromeric activity, not all are strictly necessary. Twin spot 23 originated from a symmetric event that resulted in duplication of the left arm in the green tissue and of the right arm in the white tissue (Suppl. Fig. S6). While other mechanisms may explain this outcome, the simplest interpretation is that this entailed formation of isochromosomes, indicating that the centromere overlaps the shared region, which is present in two copies.

We found four single spots and one twin spot without apparent dosage changes. In the copy-neutral single spots we detected no recombination by haplotype analysis. Their cause remains to be determined: it could be mutation or gene conversion. The twin spot event was consistent with reciprocal CO by displaying complete disomy and homozygosity. However, the crossover site was near the inferred centromere where meiotic CO are very rare. Alternatively to reciprocal CO from HR, the event could originate from two concurrent breaks followed by non-homologous end joining (NHEJ)-mediated intrachromosomal translocation. Analysis of twin and single spot formation in mutants deficient for components of different DNA repair pathways would provide additional information. Regardless of the cause of this last event, we conclude that true reciprocal CO are rare and most twins and single spots arise from genome instability leading to translocation and aneuploidy.

The observed frequency of visible spotting likely underestimates the true extent of sectoring and instability. High-magnification imaging revealed numerous microspots—green sectors comprising only one or a few cells—especially adjacent to larger sectors (Suppl. Fig. S2, S3). Equivalent white spots would be difficult to detect. Furthermore, due to layer-specific chimerism, many small green and twin spots may not be visible on the leaf surface.

Chromosome 1 does not appear to be uniquely fragile. Our data indicate that it frequently participates in translocations with other chromosomes, and we also observed coincidental instability events affecting chromosomes not linked to *CHLI1*. In addition, 6 twin spots originated from instability at Chr 23, which harbors *CHLI1-B.* This suggests that chromosome fragility and translocation potential are not restricted to the chromosomes carrying the *sulfur* locus, but are likely genome wide.

Widespread genome instability in differentiated cells could contribute to somaclonal variation—structural alterations arising during regeneration from differentiated tissues. Does the structural mutation rate vary during development? It is plausible that meristem and particularly stem cells in the L2 layer, which eventually produces the gametes, may actively protect the genome while the genome of differentiated cells may be under relaxed surveillance and repair.

Our data relied on a visual marker, chlorophyll, that could not be scored in the L1 because most epidermal cells do not contain chlorophyll. Future studies with different markers and single-cell analyses should be instrumental in quantifying structural mutation frequencies in different tissue and cell types and understanding the origins of these events.

Our results help interpret historical work on this system. Using the same *sulfur* system, Gorbunova et al. (2000) described hyper-recombination of *CHL1* after introduction of a transgenic Ac transposable element. This mutant, however, did not display hallmarks of known recombination pathway activation. Our results favor an alternative explanation considered by the authors: one or more Ac inserted into Chr 1 may induce frequent breaks near *CHLI1*, triggering translocations resembling those previously described in maize (Dooner and Belachew 1989) and in tomato (Peterson and Yoder 1993).

Why did multiple historical papers propose reciprocal CO as the cause of twin spots in plants? We believe that they were influenced by two factors. First, mitotic COs are well documented in other systems, particularly insects and fungi. Second, two carefully designed studies in plants, by Carlson in tobacco and Hirono and Redei in arabidopsis, influenced prevailing views, despite inherent limitations in their approaches. Both groups used linked heterozygous loci affecting visual traits, as molecular markers were not yet available.

Carlson employed two loci affecting leaf color, one of which was dosage sensitive. Their phenotypic similarity likely made detection and interpretation challenging. Hirono and Redei selected three loci that unbeknownst to them spanned the centromere of Chr 1: *gi* (in the upper arm, associated with plant size), *ch-1* (adjacent to the centromere, causing yellowing), and *pa* (in the lower arm, producing a dwarf phenotype). At the time, the lack of information about centromere position complicated interpretation. In addition, both *ch-1* and *pa* cause dwarfing to varying degrees, potentially confounding phenotypic classification. While these markers were the best tools available to these competent early investigators, modern molecular tools offer a chance to re-evaluate their conclusions. Our data indicate that Carlson’s inference of reciprocal crossover as the main causal mechanism of twin spots was incorrect. The conclusions of Hirono and Rédei, though more nuanced, should also be revisited.

Our study also suffers from some limitations. LOH linked to twin spots can only result from recombination occurring at the G2 phase of the cell cycle. At G1, neither reciprocal CO nor reciprocal translocation leads to observable somatic sectors because no chromatid asymmetry is generated (see Fig. S1). As such, analysis of twin spots does not allow us to test the frequency and type of recombination at G1. This is not the case for the deletions underlying single-color spots, which could arise at any cell cycle stage. This may contribute to the large disparity between the high frequency of single spots and the lower frequency of twin spots.

Theoretically, homologous recombination (HR) between chromosomes is more likely to occur at G1, when no sister chromatid is available and, for single copy DNA, the homolog is the only template for repair. However, when breaks occur in repeated DNA, multiple repeat copies may serve as templates, complicating repair outcomes. This highlights another limitation of our study: most Chr 1 breaks occurred in gene-poor, likely repeat-rich regions. Although in theory these breaks could be repaired via HR with distant repeats, repetitive DNA is typically embedded in repressive chromatin and is generally inaccessible to the recombination machinery as shown for chromoanagenesis repair (Tan et al. 2015). For this reason, we favor a model in which spontaneous double-stranded breaks in repetitive regions are repaired through non-homologous end joining (NHEJ), leading to the observed genome instability.

In conclusion, we have determined that genome instability, essentially chromosomal breaks and aneuploidy, together with the effect of meristems’ layer structure, are the major contributor to chimerism associated with LOH, including twin spots events that are often attributed to reciprocal crossovers. Leveraging these principles for the interpretation of visual markers of genomic instability should facilitate our understanding of developmental and molecular pathways that control genome function.

## Funding

Financial support for this research was provided by National Science Foundation Plant Genome Integrative Organismal Systems (IOS) Grant 1956429 (Variants and Recombinants without Meiosis) to L.C. and I.M.H.

## Methods

### Germplasms and Plant Growth Conditions

Both varieties of *Nicotiana tabacum* (Tobacco) used in this experiment were acquired from the USDA: known *sulfur* heterozygote in the background ‘John Williams Broadleaf’ (JWB) (PI551288), and strain K326 (PI552505) from which the Sierro reference genome was assembled. To confirm the *sulfur* genotype in the JWB plants, selfed offspring were grown on soil to observe segregation of the trait. Consistent with Mendelian expectations, soil-grown plants segregated 1:2 for green:yellow leaves An additional small population (n=70) were grown *in vitro* on MS media, and segregated 1:2:1 for green:yellow:white.

*Sulfur* JWB heterozygotes were outcrossed to the WT K326 plants and approximately 100,000 seeds were recovered from crosses where JWB received pollen from K326. Reciprocal crosses were also performed, and resulting fruits were harvested individually to mitigate pollen contamination. Approximately 1,200 F1 seeds from a single fruit of JWB x K326 were germinated on soil and segregated 1:1 for green:yellow. Yellow plants were transplanted to individual 5” pots approximately 3 weeks post-gemination, and green seedlings were discarded. Twin-and Single-spots were observed to originate spontaneously on new leaves throughout plant growth.

Plants were grown in greenhouse and growth chamber conditions. Greenhouse plants received no Suppl. lighting, and were kept at a daily fluctuating temperature. Two growth chambers were used: one where plants were under 16 hour days and 8 hour nights. Day conditions were 400uMol light, at 27°C, 60% relative humidity; night conditions were 22°C and 60% relative humidity. In the other growth chamber light cycles were 16 hours light (150 umol/m^2sec) at 22°C and 8 hours dark at 18 °C, kept at constant 60% humidity.. Twin spots and single spots arose on plants in all growth conditions, and tissue was collected from all three environments.

Spot counts were performed on plants under constant 23°C and 75% humidity, with 16 hour days, 8 hour nights, and 150 umol/m^2sec of light.

### Spot Observations

Observations of mosaic sectors were done 8 weeks after planting on JWB x K326 sulfur heterozygotes. Plants with at least 7 leaves were counted, with the cotyledons, the two oldest leaves, and the 3 newest leaves still in development being excluded. Spots were counted visually, and a ruler with 1 mm markings was used to measure spot size. All spots with a cross-section >1mm in any direction were counted. Spots with a cross-section >5mm were noted. Spots were counted for each leaf individually, and summed to provide the total number for each plant. Spots were also grouped into one of three categories: green single spots were sectors of tissue which were darker than the background tissue, white single spots were sectors lighter than the background tissue, and twin spots were sectors where there was adjacent green and white tissue.

### Tissue Collection

Tissue used for sequencing was collected from twin- and single-spots occurring on leaves of F1 JWB x K326 plants. For each spot, tissue of each color-type was excised using a sterile blade, with a margin left at the border to mitigate contamination from adjacent tissue. In addition to the green and white tissue, yellow tissue adjacent to the spot was collected. Harvested tissue was flash-frozen in liquid nitrogen and stored at −80°C until DNA extraction.

Tissue used for RNA extraction and sequencing was collected from the fifth leaf, sized approximately 2.5 cm, from each of six phenotypically yellow su/+ and six green WT seedlings in a homozygous JWB background. Due to the delayed growth of *su/*+ relative to WT siblings, seeds segregating 1:2 WT:*su/+* on soil were planted in successive batches in order to sample leaves of similar size and developmental stage. Leaf tissue of both genotypes was collected on July 12, 2022 from 10:45-10:55 am and immediately snap-frozen in liquid nitrogen.

### RNA Preparation and Sequencing

Total RNA was extracted from each frozen leaf separately using TRIzol and the Qiagen RNeasy plant mini kit protocol. Tissue was ground to a fine powder in a pre-chilled mortar and pestle, and 1 mL of TRIzol was added directly to the mortar and homogenized further. Samples were then brought to room temperature, transferred to RNAse-free 1.7 mL tubes, and incubated for five minutes at room temperature. Subsequent RNA extraction steps were performed using a Qiagen RNeasy plant mini kit according to manufacturer protocol. RNA sample integrity was assessed by gel electrophoresis.

Sequencing libraries were prepared using a KAPA Hyper Prep mRNA kit (catalog no. 08098123702) protocol and reagents. Sample concentrations were measured using a QuBit 2.0 fluorometer, pooled in equimolar amounts and sequenced using an Element Biosciences AVITI sequencer at the UC Davis DNA Technologies Core.

### DNA Preparation and Sequencing

DNA extraction was performed using a CTAB extraction method. Frozen leaf tissue was ground in a chilled 1.5mL tube using a chilled plastic homogenizer for approximately 20 seconds until a fine powder was formed. Pre-warmed CTAB buffer was added and samples were incubated for 30 minutes in a 60°C water bath with intermittent vortexing. DNA was extracted once by adding 24:1 chloroform:isoamyl alcohol and centrifuging at 6,000x g for 10 minutes. The clear aqueous phase was extracted and nucleic acids were precipitated using cold isopropanol. Samples were centrifuged, the supernatant was discarded, and 70% ethanol was added to precipitate the DNA once more. After centrifuging, the supernatant was discarded and pelleted DNA was allowed to air dry briefly at room temperature before resuspension in 10mM Tris-HCl (pH 8.0). RNAse was added, and samples were incubated at 37°C for 30 minutes. Samples were stored at −20°C.

To prepare for sequencing, DNA samples were sonicated to a fragment size of 300-500 bp using a Covaris sonicator (175W peak power, 10% duty factor, 200 cycles/burst, 90s treatment time). Fragmented DNA was size-selected by addition of equal volume of KAPA Pure magnetic beads, and elution in 10mM Tris-HCl (pH 8.0). Genomic libraries were prepared using the KAPA DNA HyperPrep protocol and reagents. Sample concentration was measured using a QuBit 2.0 fluorometer and standardized to 5ng/uL. Normalized samples were pooled and sequenced to give approximately 10 million reads per sample using Element Biosciences AVITI sequencing system and Illumina NovaSeq available through the UC Davis DNA Technologies Core.

Sequencing of the *sulfur* SNP was done using the PCR sequencing services offered by Plasmidsaurus. Primers AP8 and AP9 (table x) were used to amplify a 2kb region surrounding the identified SNP. The PCR was set up using Takara ExTaq DNA polymerase and reagents, a high-fidelity polymerase suitable for long-range PCR. PVP was added to PCR buffers at a final concentration of 1.5%. The reaction was annealed at 63°C and a total of 35 PCR cycles were performed. Products were run on a 1% agarose gel in a TAE buffer at 120V for 20 minutes to confirm adequate amplification prior to sending for Plasmidsaurus sequencing.

### Sequencing Data Analysis

Sequencing data from AVITI was demultiplexed using a custom python script (https://comailab.org/data-and-method/barcoded-data-preparation-tools-documentation/). Reads were aligned to the K326 (Sierro et al) reference genome using BWA, and the resulting files (.sam files) were used for dosage analysis. A custom script (https://github.com/ComaiLab/bin-by-sam) was used to generate dosage plots, where first mapped reads were binned into 1 Mb consecutive, non-overlapping bins. Next, relative coverage was calculated by normalizing read number per bin to a control sample. In this case, the control used was the yellow tissue from green single spot 3, as it had the best read coverage among the first set of sequenced spots. The resulting data was visualized using Rstudio and plots were visually inspected for dosage changes. For spots with breakages, the break locus on each chromosome was estimated within a 2 Mb range based on the bins flanking a change in dosage.

To genotype the tissue samples, we first identified SNPs between the reference genome (K326) and sulfur JWB, by pooling all sequencing data from all tissue sampled from F1 JWB x K326 into a single file. Heterozygous positions with coverage between 40 and 100 and the two alleles each represented between 40% and 60% of the basecalls were categorized as SNPs between the reference K326 and JWB (alternative allele). Next, each sample was processed independently. For each consecutive non-overlapping 2.5 Mb bin of the reference genome, the percentage of reads supporting a reference (K326) allele based on those SNP positions was recorded and expressed as % Reference (WT) allele.

RNA-seq reads were aligned to the Edwards reference genome (Edwards et al. 2017) using HISAT2 (Kim et al. 2019), PCR duplicates removed with Picard MarkDuplicates (version 2.18), and converted to binary format with samtools. The putative *sulfur* locus was identified through BLASTN alignment of the transposon tag sequence from Fitzmaurice et al (1999). Read alignments at the putative *sulfur* locus were visualized with IGV.

## Suppl

Suppl. online Data

Online Suppl. data table 1. Contains summary of spot analysis. https://figshare.com/s/bfad31f2a151e5acdf8d

## Supplemental Figures

**Supplemental Figure S1.**
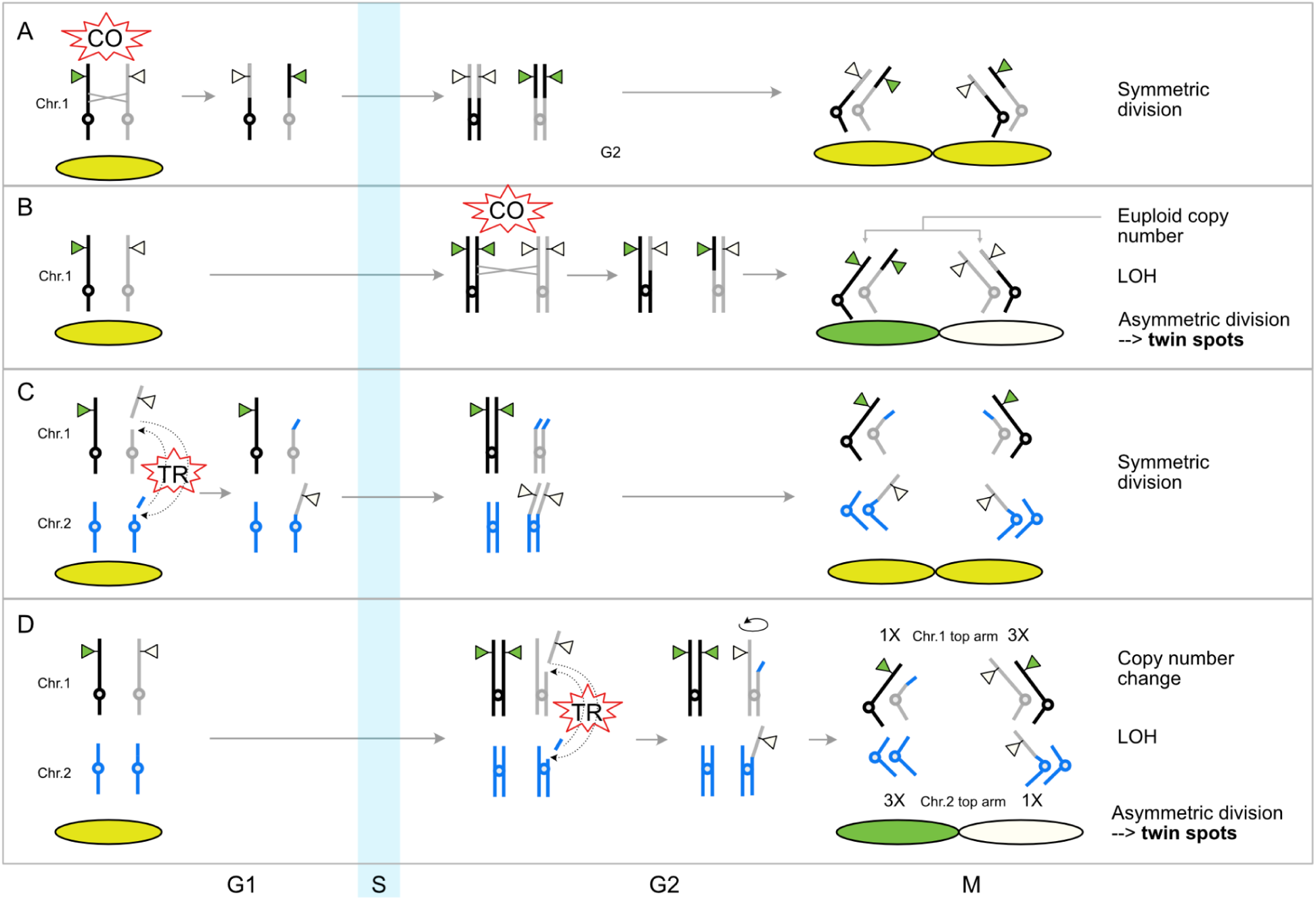
Timing of recombination events and twin spots formation. The green and white triangle represent, respectively, + (WT) and *su* (mutant) alleles of the CHL1 gene. The *+/su* F1 hybrid used in this study is represented by the black and grey haplotypes of Chr 1 carrying, respectively, + and *su* alleles. The drawing illustrates how the outcome of recombination events depends on cell cycle timing. A. Reciprocal homologous crossover (CO) at G1 phase between the two homologs of Chr 1 resulting in recombinant chromosomes. Because recombination occurs before replication, the CO does not produce asymmetric chromatids and has no somatic impact. B. The CO takes place at G2 affecting single chromatids and creating an asymmetric allelic conformation. At anaphase of mitosis, there is 0.5 probability of an asymmetric division and LOH resulting in twin spots. C, D. Similarly to A, B the effect of a translocation depends on cell phase timing. In C, a reciprocal translocation (TR) that occurred at G1 has no somatic effect. In D, however, the TR generated asymmetric chromatids and has 0.5 probability of causing LOH and twin spots.

**Supplemental Figure S2.**
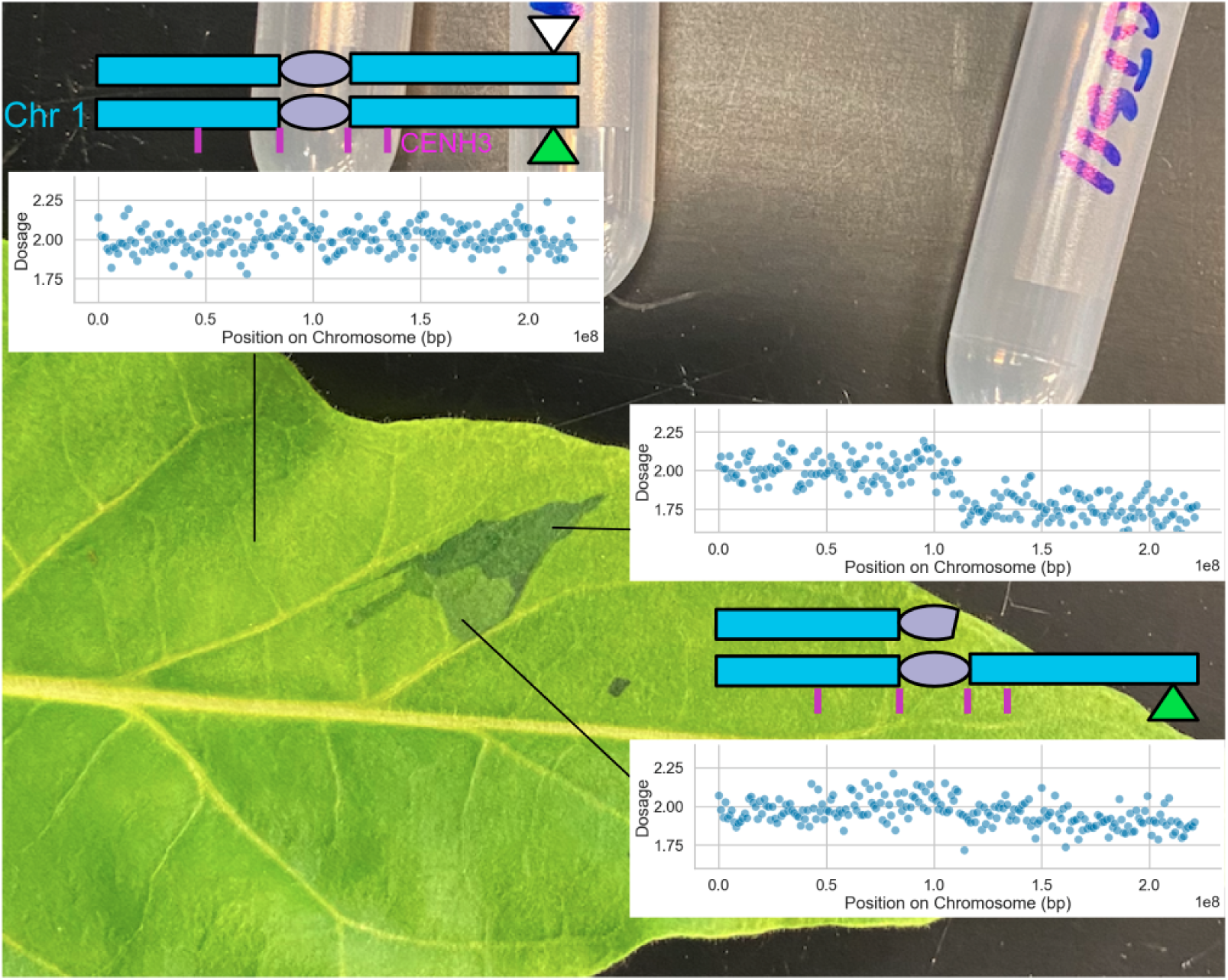
Layer chimerism appears as a false twin spot. *N. tabacum sulfur* heterozygote leaf. This single spot was originally mistaken for a twin spot. After the dosage analysis, it was apparent that the “white” and green portions of the twin were caused by the same lesion: deletion of the right chromosomal arm containing the *su* haplotype. The “white” spot was instead a green spot of a lighter shade than its apparent twin. The twinning effect was the result of differential layer invasions. There were fewer mutant cells in the light green spot and the dosage shift was accordingly smaller than the shift in the green spot.

**Supplemental Fig. S3.**
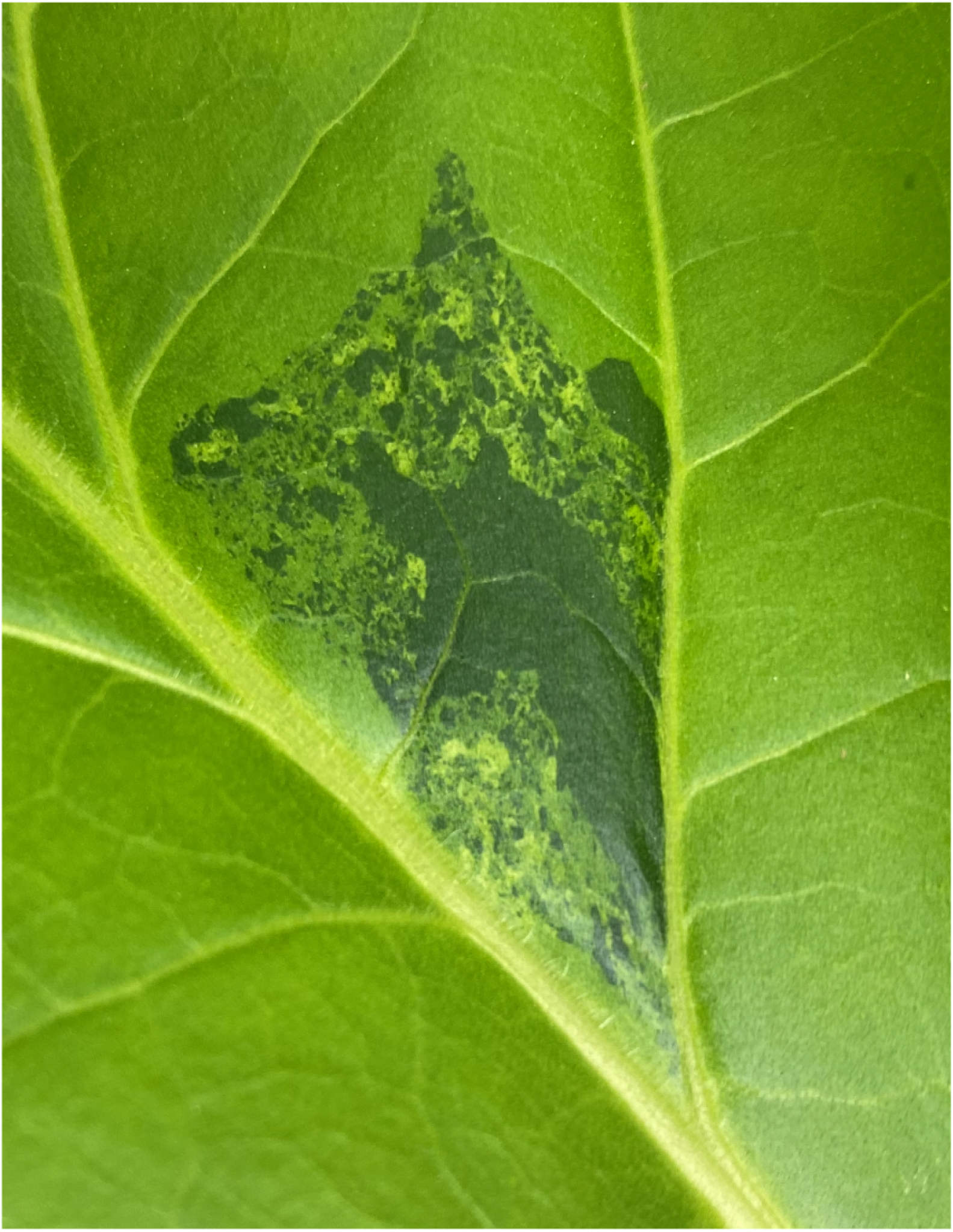
Composite spotting indicating genome instability. *N. tabacum sulfur* heterozygote leaf.

**Supplemental Figure S4.**
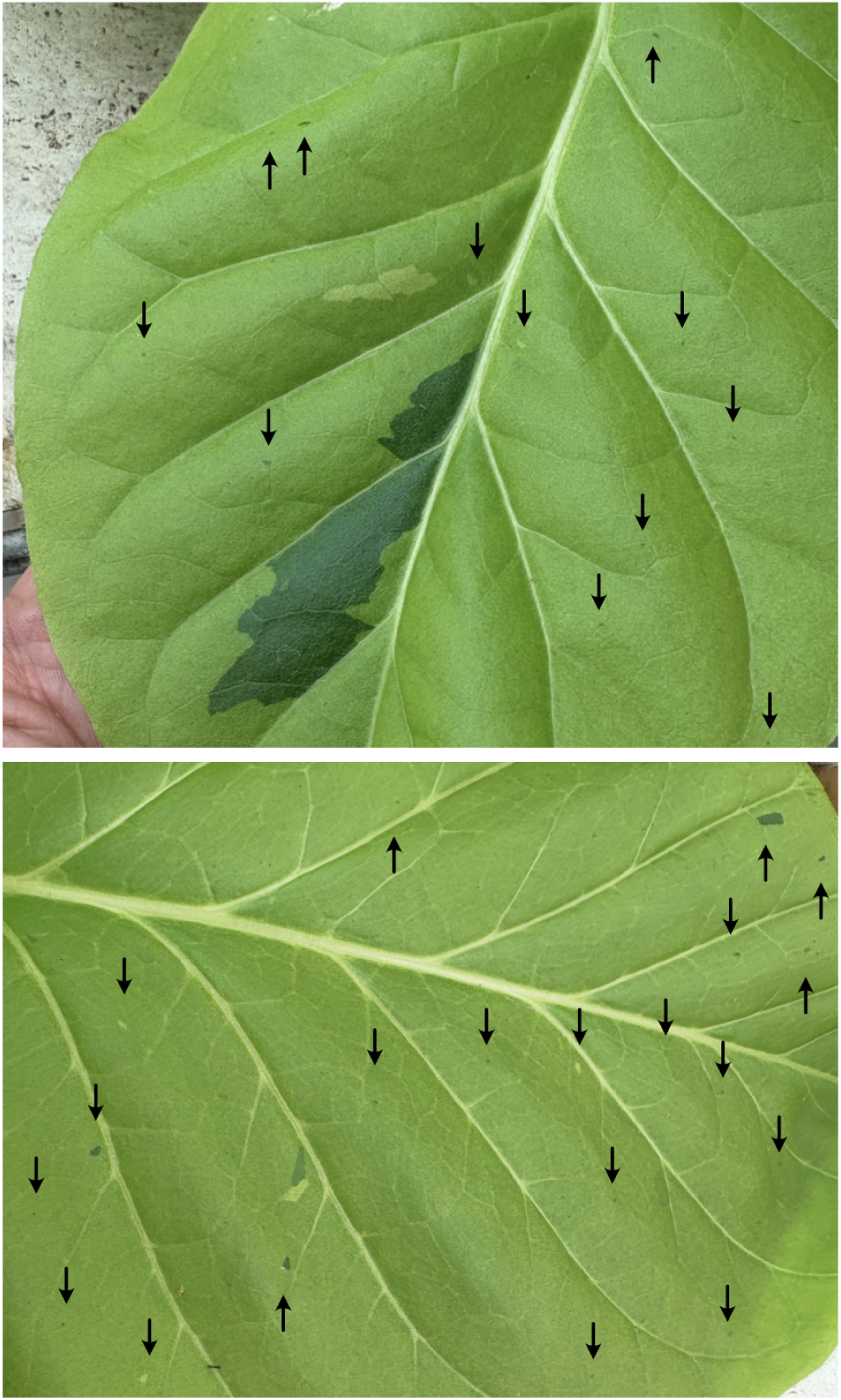
Spot clusters. *N. tabacum sulfur* heterozygote leaves. In each image, spots of varying dimension (arrows) accompany one or two larger spots. Instability, therefore, was associated with development of each leaf indicating that either a genetic or epigenetic property was associated with to spot formation.

**Supplemental Fig. S5.**
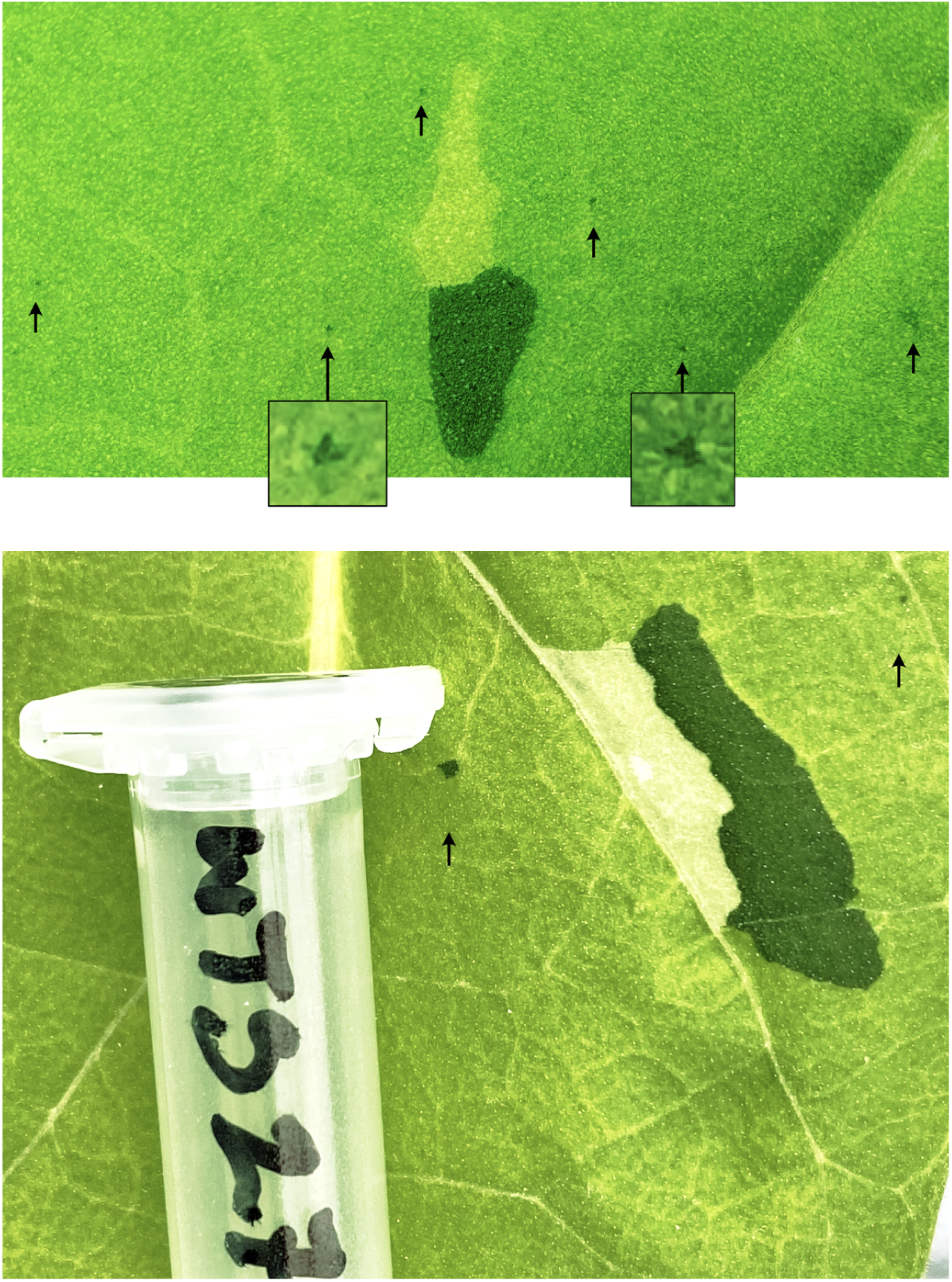
Twin spots and associated microspots. *N. tabacum sulfur* heterozygote leaves. The arrows point to small green spots, two of which are enlarged. Top: Twin spots formed on the F1 hybrid su/+ during May 2025. The DNA was not sampled. Bottom: Twin spot 27. Formed during May 2023 and sampled for DNA analysis. The effect of layer penetration by chimeric cells on the color shade is distinct. In twin spot 27, the colors are particularly vivid suggesting that multiple and superficial cell layers were affected. The Variant Allele Frequency for the corresponding LOH was 0.65 - 0.35 (1-0 would be expected) indicating that some layers maintained the ancestral *su/+* genotype. These could be, for example, upper and lower epidermis and mesophyll layers in the adaxial side.

**Supplemental Fig. S6.**
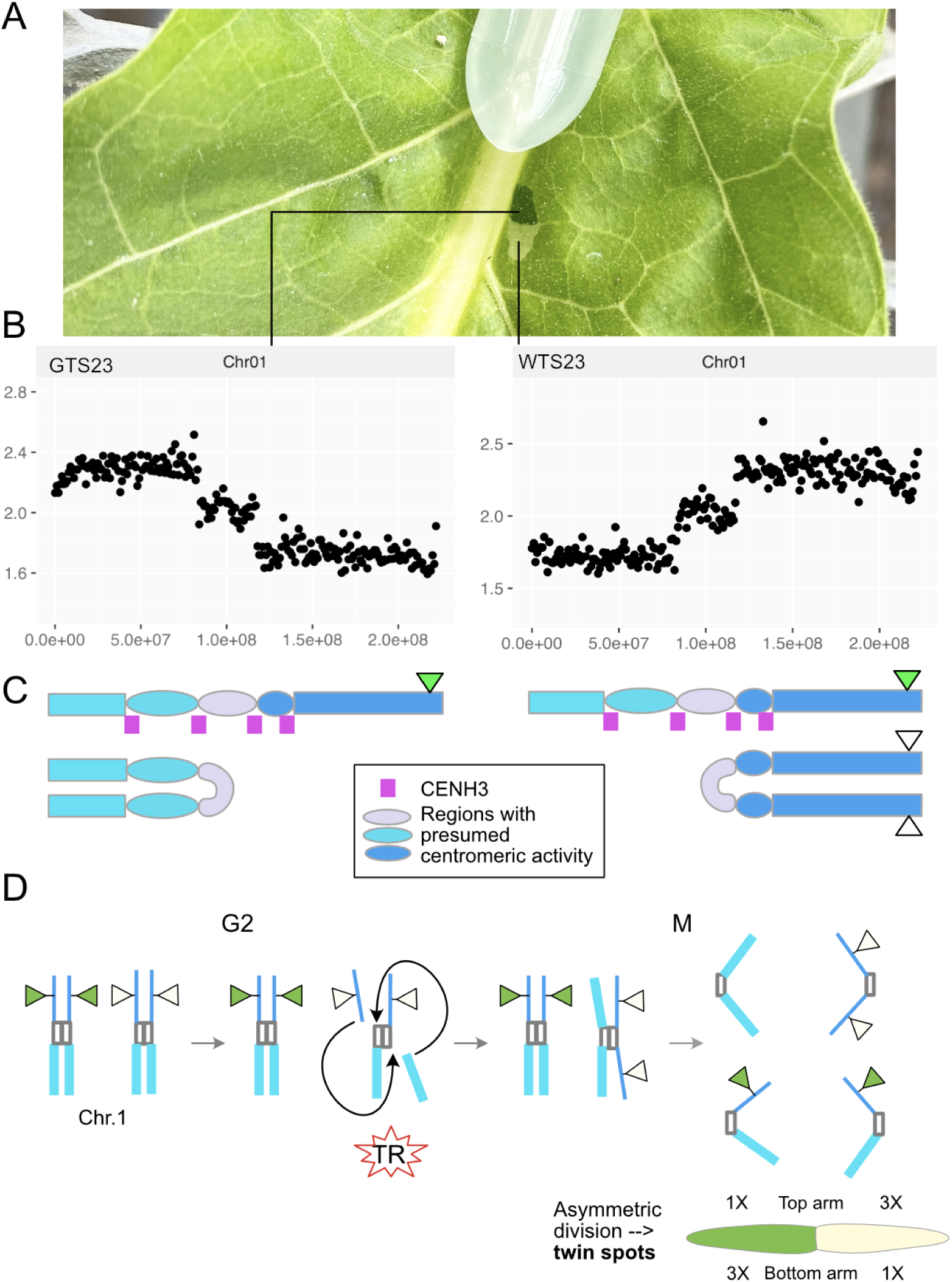
Analysis of internal translocations associated with twin spot 23. Genome wide dosage analysis revealed that the only genomic alteration affected Chr 1. (A) Twin spot 23 formed on the leaf of *N. tabacum sulfur* heterozygote. (B) The dosage of green and white spot DNA displays, respectively and symmetrically, 3:1 and 1:3 copies of the left and right arm. The middle region is present in 2 copies. (C) We interpret this outcome as the result of symmetrical isochromosome formation. This hypothetical rearrangement would place the centromere in the indicated interval. (D) Model for symmetric formation of two isochromosomes with overlapping centromeric regions. The published CENH3 profile for the region is displayed in magenta ticks (Sierro et al. 2024; Zan et al. 2025) illustrating the coincidence of the two central CENH3 signals, the detected breaks, and the predicted junctions. TR: translocation.

**Supplemental Figure S7.**
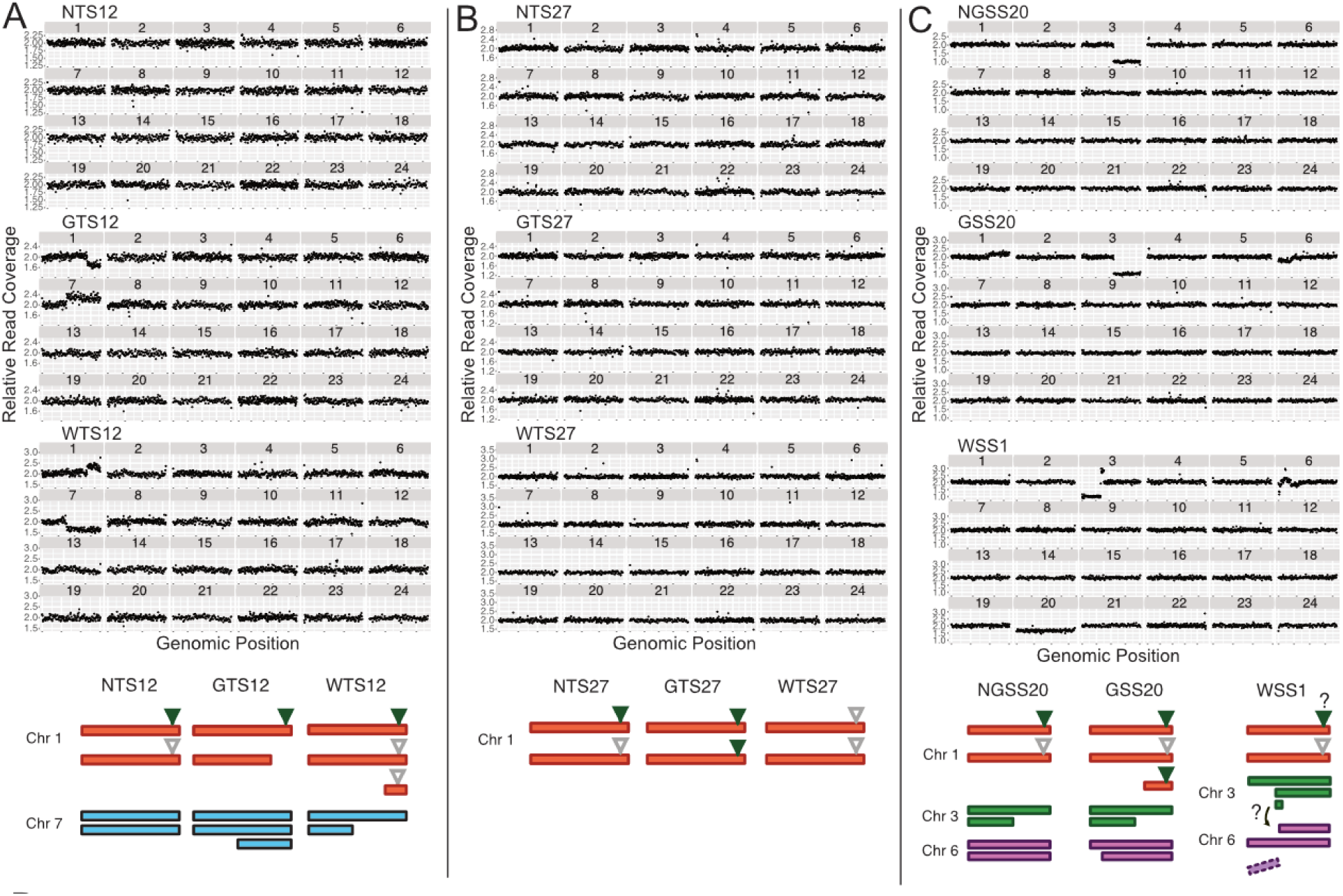
Relative dosage analysis of Twin spot 12, Twin spot 27, Green single spot 20 and White single spot 1. The graphs represent the genomic dosage profile of each sampled spot. NTS: control leaf region displaying the yellow trait and adjacent to the sampled twin spot. GTS: Green spot of twins. WTS: White spot of twins. NGSS: control leaf region displaying the yellow trait and adjacent to the sampled green spot. GSS: green single spot. WSS: white single spot. The lower schematics represent the inferred karyotype.

**Supplemental Figure 8.**
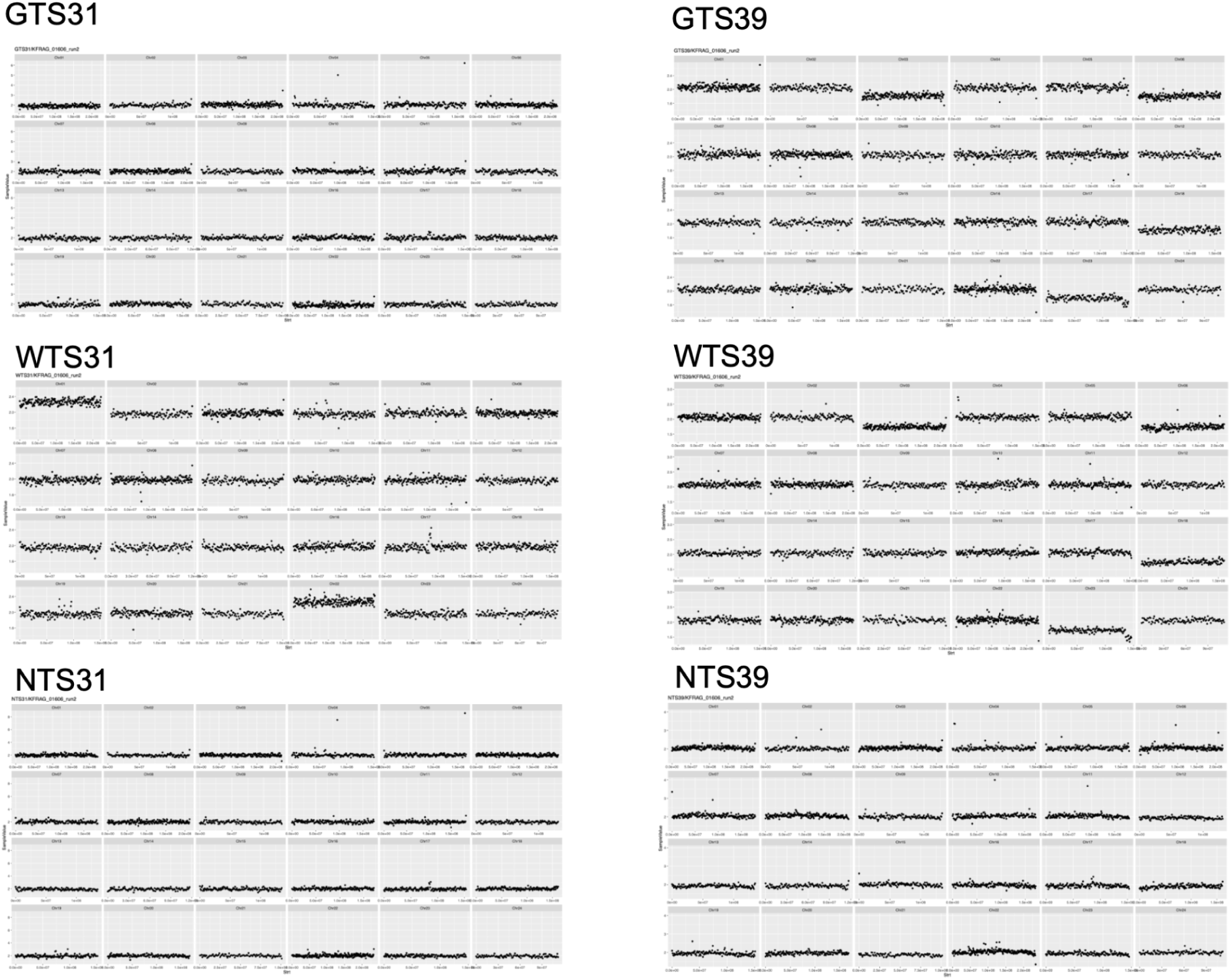
Genomic dosage profiles for twin spot 31 and twin spot 39 reveal aneuploidy.

**Supplemental Figure S9.**
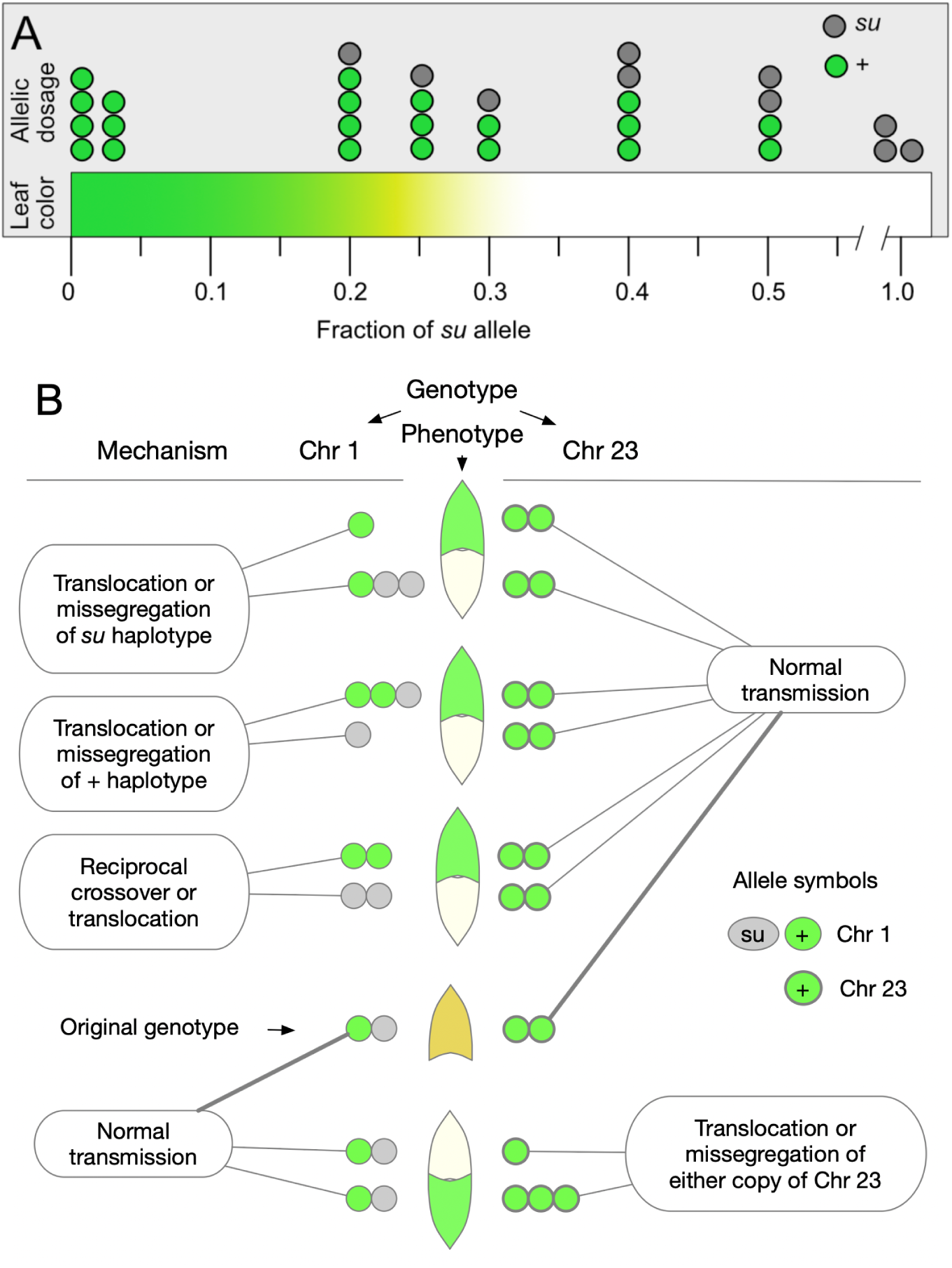
Schematic representation of allelic contributions from the two active Mg Chelatase loci. (A) Effect of allelic dosage on leaf color. (B) Mitotic inheritance and interaction of CHLI1-A on Chr 1 and CHLI1-B on Chr 23 genotypes in the formation of twin spots.

---

**Supplemental Table S1.** Count of total spots >1mm for 25 plants at 8 weeks post-planting. Any spots greater than 5mm in diameter were noted as well as those larger than 1cm (columns 6 & 7). Notably, all plants produced multiple green single spots, at least one white spot, and all but two plants produced at least one twin spot. Cotyledons, the two oldest true leaves, and developing leaves were not scored.

| Plant | Number of Leaves Scored | GSS >1mm | WSS >1mm | TS >1mm | spots >5mm | spots >1cm |
| --- | --- | --- | --- | --- | --- | --- |
| ka 366 4 | 6 | 61 | 14 | 3 | 2gss |  |
| ka 366 5 | 4 | 69 | 15 | 3 | 1ts 1gss |  |
| ka 369 5 | 5 | 52 | 5 | 2 | 1ts 1wss 2gss |  |
| ka 367 7 | 4 | 61 | 12 | 3 | 1ts 1gss |  |
| ka 366 11 | 7 | 62 | 21 | 7 | 1ts 1wss 1gss | 1gss |
| ka 368 4 | 4 | 31 | 8 | 0 |  | 1gss |
| ka 368 3 | 6 | 40 | 11 | 1 |  | 2gss |
| ka 367 5 | 5 | 75 | 20 | 5 | 1ts 1gss 1wss |  |
| ka 369 4 | 4 | 36 | 5 | 2 | 1ts |  |
| ka 367 4 | 4 | 16 | 2 | 2 | 1wss 1ts |  |
| ka 368 2 | 4 | 32 | 15 | 0 | 2gss |  |
| ka 368 5 | 6 | 75 | 20 | 5 | 2gss |  |
| ka 368 1 | 3 | 25 | 7 | 1 |  |  |
| ka 367 3 | 4 | 59 | 16 | 3 | 2gss |  |
| ka 367 2 | 8 | 70 | 18 | 6 | 1ts 5gss |  |
| ka 369 3 | 5 | 32 | 5 | 6 | 2ts |  |
| ka 367 13 | 5 | 28 | 5 | 1 | 1wss 1gss |  |
| ka 366 3 | 6 | 60 | 17 | 3 | 1wss |  |
| ka 369 2 | 6 | 67 | 23 | 4 | 2gss | 1gss |
| ka 369 1 | 5 | 36 | 18 | 4 | 1ts 1gss |  |
| ka 367 1 | 7 | 47 | 6 | 1 | 4gss |  |
| ka 366 7 | 5 | 26 | 9 | 1 |  |  |
| ka 366 6 | 5 | 34 | 8 | 3 | 1wss | 1wss 1gss |
| ka 366 9 | 5 | 47 | 17 | 4 | 1gss |  |
| ka 366 8 | 5 | 53 | 9 | 3 |  |  |
| ka 367 6 | 3 | 16 | 1 | 1 |  |  |
| Totals | 131 leaves | 1210 | 307 | 74 | 28gss, 11ts, 7wss | 6gss, 1wss |

**Supplemental Table S2.** Regions of chromosomes involved in translocation and data on chromosome 23 break sites. For chromosome 23, all the breaks spanned from locus 1 to the site indicated in the table. For all these chromosomes, the breaksite was compared with predicated CENH3 binding and it was indicated if the break region fell within a peak. Only 4 spots had the nonhomologous chromosome translocation occurring in a CENH3 binding site.

| Spot Number | Other Chr Altered | Approx Breakage Region | In Peak (cenH3 bins with relative coverage >2) |
| --- | --- | --- | --- |
| TS1 | 7 | 1 to 1.1-1.3e7 | No |
| TS2 | 13 | 1.07-1.09e8 to end | No |
| TS5 | 17 | 1 to 5e6-7e6 | No |
| TS7 | 12 | 1 to 6e6-8e6 | No |
| TS8 | 13 | 1.07-1.09e8 to end | No |
| TS9 | 8 | 1.2-1.22e8 to end | No |
|  | 8 | 8.7-8.9e7 to end | No |
| TS10 | 10 | 1 to 4.0 - 4.2e7 | Yes |
|  | 10 | 1 to 4.2-4.4e7 | Yes |
| TS12 | 7 | 7.5 - 7.7e7 to end | No |
| TS18 | 22 | 4.9-5.1e7 | Yes |
| TS23 | 1 | 1 to 8.4-8.6e7 | No |
| TS30 | 6 | 1 to 7.6-7.8e7 | Yes |
| TS32 | 2 | 5.8-6.0e7 to end | No |
| TS34 | 5 | 1.08-1.16e8 to end | No |
| TS36 | 19 | 9.6e7 - 1e8 to end | Yes |
| TS14 | 14 | 1 to 2.9-3.1e7 | No |
|  | 12 | 1 to 3.3-3.5e7 | No |
| GSS1 | 13 | 1.07-1.09e8 to end | No |
| GSS3 | 13 | 1 to 1.3-1.5e7 | No |
| GSS20 | 3, 6 | (chr6) 1 to 5.0e7-5.2e7 | No |
| GSS32 | 2 | 1 to 4.6-4.8e7 | No |
| WSS1 | 3, 6, 20 |  |  |
| TS8 | 23 | 2.8-3.0e7 | No |
| TS9 | 23 | 3.0-3.2e7 | No |
|  | 23 | 3.6-3.8e7 | No |
| TS18 | 23 | 4.9-5.1e7 | No |
| TS34 | 23 | 4.5-4.7e7 | No |
| WSS3 | 23 | 2.1-2.3e7 | No |
| WSS17 | 23 | 4.6-4.8e7 | No |

**Supplemental Table S3.** Phenotypes of pseudo-backcross and F2 populations segregating *su/+*.

| Population | Condition | Green | Yellow | White |
| --- | --- | --- | --- | --- |
| F1 (*su/+* X K326) | MS agar 1% sucrose | 37 | 41 | 0 |

## Notes

https://figshare.com/s/bfad31f2a151e5acdf8d

### Competing Interest Statement

The authors have declared no competing interest.

